# Coxsackievirus infection induces direct pancreatic β-cell killing but poor anti-viral CD8+ T-cell responses

**DOI:** 10.1101/2023.08.19.553954

**Authors:** Federica Vecchio, Alexia Carré, Daniil Korenkov, Zhicheng Zhou, Paola Apaolaza, Soile Tuomela, Orlando Burgos-Morales, Isaac Snowhite, Javier Perez-Hernandez, Barbara Brandao, Georgia Afonso, Clémentine Halliez, John Kaddis, Sally C. Kent, Maki Nakayama, Sarah J. Richardson, Joelle Vinh, Yann Verdier, Jutta Laiho, Raphael Scharfmann, Michele Solimena, Zuzana Marinicova, Elise Bismuth, Nadine Lucidarme, Janine Sanchez, Carmen Bustamante, Patricia Gomez, Soren Buus, the nPOD-Virus Working Group, Sylvaine You, Alberto Pugliese, Heikki Hyoty, Teresa Rodriguez-Calvo, Malin Flodstrom-Tullberg, Roberto Mallone

**Affiliations:** Université Paris Cité, Institut Cochin, CNRS, INSERM, Paris, France; Institute of Diabetes Research, Helmholtz Zentrum München, German Research Center for Environmental Health, Munich-Neuherberg, Germany; German Center for Diabetes Research (DZD), Neuherberg, Germany; Center for Infectious Medicine, Department of medicine Huddinge, Karolinska Institutet, Karolinska University Hospital Huddinge, Stockholm, Sweden; Diabetes Research Institute, Leonard Miller School of Medicine, University of Miami, FL, USA; Department of Diabetes Immunology, Arthur Riggs Diabetes and Metabolism Research Institute, Beckman Research Institute, City of Hope, Duarte, CA, USA; Department of Diabetes and Cancer Discovery Science, Arthur Riggs Diabetes and Metabolism Research Institute, Beckman Research Institute, City of Hope, Duarte, CA, USA; University of Massachusetts Medical Chan School, Diabetes Center of Excellence, Department of Medicine, Worcester, MA, USA; Barbara Davis Center for Diabetes, University of Colorado School of Medicine, Aurora, CO, USA; Islet Biology Exeter (IBEx), Exeter Centre of Excellence for Diabetes Research (EXCEED), Department of Clinical and Biomedical Sciences, University of Exeter Medical School, Exeter, UK; ESPCI Paris, PSL University, Spectrométrie de Masse Biologique et Protéomique, CNRS UMR8249, Paris, France; Tampere University, Faculty of Medicine and Health Technology and Fimlab Laboratories, Tampere, Finland; Paul Langerhans Institute, Technical University Dresden, Germany; Assistance Publique Hôpitaux de Paris, Service d’Endocrinologie Pédiatrique, Robert Debré Hospital, Paris, France; Assistance Publique Hôpitaux de Paris, Service de Pédiatrie, Jean Verdier Hospital, Bondy, France; Department of Pediatrics, Division of pediatric Endocrinology, Leonard Miller School of Medicine, University of Miami, FL, USA; Panum Institute, Department of International Health, Immunology and Microbiology, Copenhagen, Denmark; Indiana Biosciences Research Institute, Indianapolis, IN, USA; Assistance Publique Hôpitaux de Paris, Service de Diabétologie et Immunologie Clinique, Cochin Hospital, Paris, France

**Keywords:** cytotoxic T lymphocytes, Enterovirus, epitopes, HLA, immunopeptidome, glutamic acid decarboxylase, mimicry, type 1 diabetes, vaccination

## Abstract

Coxsackievirus B (CVB) infection of pancreatic β cells is associated with β-cell autoimmunity. We investigated how CVB impacts human β cells and anti-CVB T-cell responses. β cells were efficiently infected by CVB *in vitro*, downregulated HLA Class I and presented few, selected HLA-bound viral peptides. Circulating CD8^+^ T cells from CVB-seropositive individuals recognized only a fraction of these peptides, and only another sub-fraction was targeted by effector/memory T cells that expressed the exhaustion marker PD-1. T cells recognizing a CVB epitope cross-reacted with the β-cell antigen GAD. Infected β cells, which formed filopodia to propagate infection, were more efficiently killed by CVB than by CVB-reactive T cells. Thus, our *in-vitro* and *ex-vivo* data highlight limited T-cell responses to CVB, supporting the rationale for CVB vaccination trials for type 1 diabetes prevention. CD8^+^ T cells recognizing structural and non-structural CVB epitopes provide biomarkers to differentially follow response to infection and vaccination.

## Introduction

Environmental factors weigh heavier than genetic predisposition in the pathogenesis of type 1 diabetes (T1D)^1^, but remain elusive. Moreover, the increasing prevalence of neutral and protective HLA Class II haplotypes in the T1D population^2^ and the parallel increase in disease incidence^3^ suggest that environmental triggers are gaining importance. Such triggers are likely to exert their role early in life, as most children who subsequently develop T1D seroconvert for islet auto-antibodies (aAbs) before 2 years of age^4,5^.

Infections by Enteroviruses such as Coxsackieviruses B (CVBs) are suspected triggers of T1D^1^. Key features relevant to this hypothesis are the virus capacity to infect pancreatic β cells, their high prevalence (>95% of the population being seropositive) and high incidence in infants and toddlers, and their oro-fecal transmission. CVBs are held as plausible candidates based on association studies in prospective cohorts of genetically at-risk children. These studies documented a temporal correlation between serological evidence of infection by some CVB serotypes and aAb seroconversion. Using fecal metagenome sequencing, the TEDDY study^6^ documented that the predisposing effect of CVB infections is on islet autoimmunity (i.e. aAb seroconversion) rather than on its subsequent progression to clinical T1D; and that such predisposing effect is linked to prolonged rather than to short, independent CVB infections. These prolonged infections, identified by the protracted shedding of the same CVB serotype in sequential stool samples, are indicative of viral persistence^7^, which may underlie a less effective anti-viral immunity.

In parallel, spatial associations have been highlighted in histopathological studies on pancreas specimens^8–10^. Positive staining for enteroviral VP1 protein was more prevalent in T1D than in non-diabetic donors and associated with the two histopathology hallmarks of T1D, namely immune infiltration and HLA Class I (HLA-I) hyper-expression. Only a minority of islets with residual β cells were VP1^+^, which may reflect low-grade infections persisting years after the initial CVB exposure.

Whether these associations underlie a cause-effect relationship remains unsettled. T1D primary prevention trials based on a multivalent inactivated CVB vaccine^11^ are being considered^10^. Such trials are needed to directly test whether preventing CVB infection affords protection from islet autoimmunity and subsequent T1D, which may support the pathogenic role of CVB. Several knowledge gaps however exist about the natural immune response against CVB^1^. Available data is limited to serological studies, with one report suggesting that children who subsequently develop early anti-insulin aAbs lack anti-VP1 neutralizing antibodies (Abs)^12^. In line with the TEDDY study^6^, this could predispose to infections at higher viral loads, which may favor viremia and dissemination to the pancreas.

CD8^+^ T cells play a key role in the clearance of viral infections through cytotoxic lysis of infected cells, which process viral proteins and present peptides on surface HLA-I for T-cell recognition^13^. Against this background, it is unknown whether islet destruction and autoimmune initiation reflect a direct, primary β-cytolytic effect of viral infection or an indirect, secondary effect of anti-viral T cells against infected β cells^1^. The lack of knowledge about the CVB peptides processed and presented by infected β cells and recognized by CD8^+^ T cells hampers our possibility to understand this process and to follow response to infection and, eventually, vaccination. We aimed to fill these gaps by identifying the HLA-I-bound viral peptides presented by infected β cells, and by using these peptides to track CVB-reactive CD8^+^ T cells and their cytotoxic activity against β cells.

## Results

### CVB-infected β cells downregulate surface HLA-I expression and present few selected HLA-I-bound viral peptides

To identify the CVB peptides naturally processed and presented by β cells, we infected the ECN90 β-cell line^14,15^ (HLA-A*02:01/A*03:01^+^; HLA-A2/A3 from hereon), which expresses the Coxsackievirus/Adenovirus receptor (CAR) and decay-accelerating factor (DAF)^16^ required for CVB entry (not shown). *In-vitro* infection with either CVB3 or CVB1 was highly efficient, leading to expression of VP1 and double-stranded (ds)RNA in >70% of cells within 4 h (Fig. 1A). Increased β-cell death was apparent at 8 h. For immunopeptidomics experiments, we selected the 6-h time point, which corresponds to the infection plateau preserving viability. Notably, infected cells down-regulated HLA-I expression (Fig. 1B-C).

**Figure 1.**
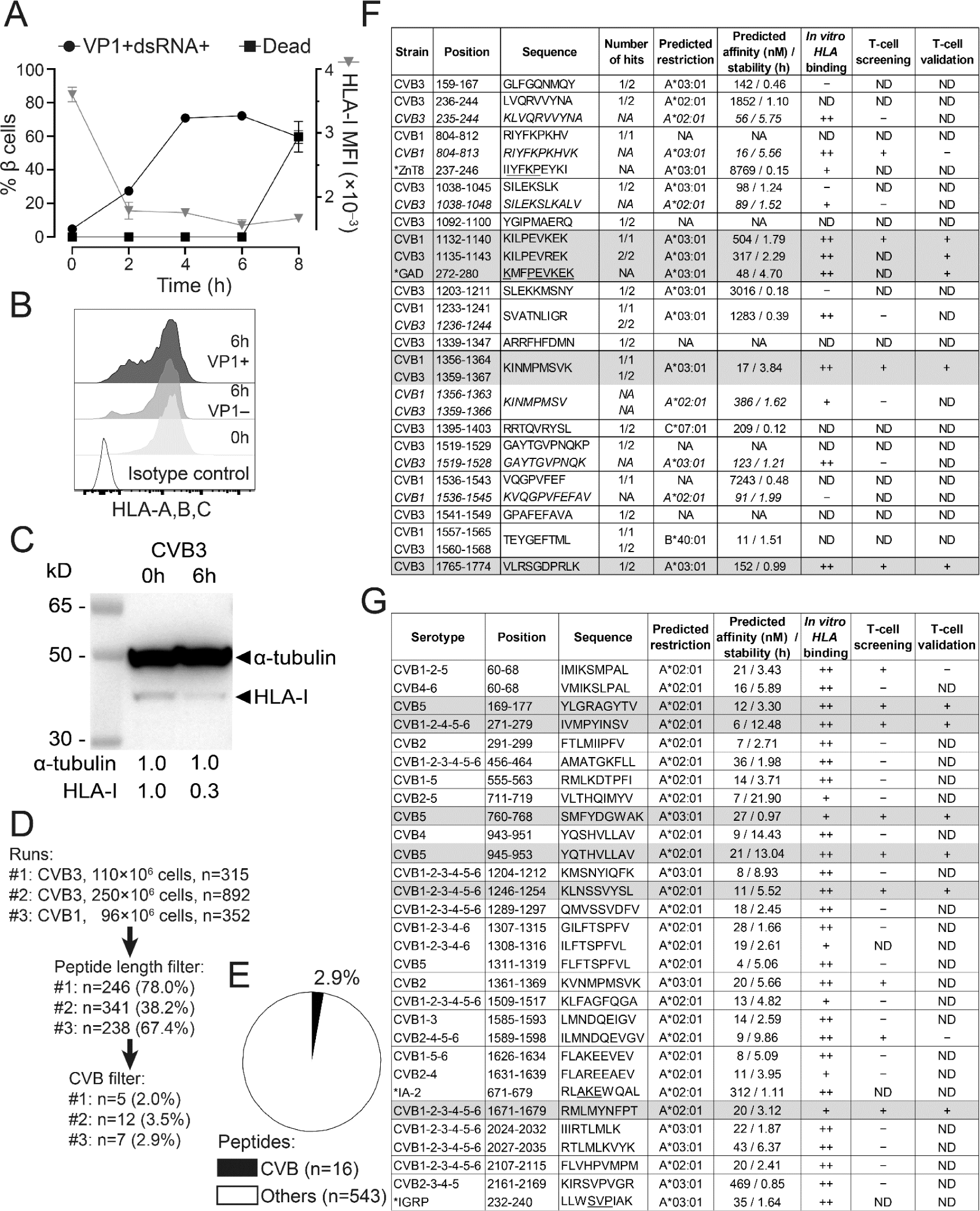
CVB-infected β cells downregulate surface HLA-I expression and present few selected HLA-I-bound viral peptides. **A.** ECN90 β cells were infected *in vitro* with CVB3 at 300 MOI. The kinetics of expression of the viral markers VP1 and dsRNA, of cell death (left y-axis) and of HLA-I median fluorescence intensity (MFI; right y-axis) was measured by flow cytometry. **B-C.** HLA-I expression measured by flow cytometry (B) and Western blot (C) on ECN90 β cells 6 h post-infection. **D.** The 6-h time point was chosen and 3 HLA-I peptidomics runs performed. Cells were lysed and HLA-I-bound peptides eluted and sequenced by MS. Sequences were filtered for peptide length (8-12 aa) and for matches with the translated RNA sequences of the CVB strains used. The number of peptides retained at each filtering step are shown (percent of peptides out of those from the previous step in parentheses). **E.** Percent CVB peptides out of total 8-12mer peptides eluted. **F.** CVB peptides retrieved from HLA peptidomics experiments and length variants of the core sequences identified (shown in italics). The number of hits out of the 3 peptidomics runs performed (Fig. 1D) are listed for each peptide, and their mapping in the CVB polyprotein sequence is detailed in <u>Fig. S1</u>. A ZnT8_237-246_ peptide homologous to CVB1_804-812_ (identified by Blast search) and a GAD_272-280_ peptide homologous to the CVB1_1132-1140_/CVB3_1135-1143_ sequence retrieved from our previous datasets^15^ are indicated with an asterisk (conserved aa underlined). HLA-I restrictions were assigned using NetMHCStabpan 1.0, and predicted affinity and stability values are listed. Subsequent columns list *in-vitro* HLA binding results (detailed in <u>Fig. S2</u>) and peptides identified as recognized by CD8^+^ T cells in the screening (Fig. 2) and validation runs (Fig. 3). Peptides eventually validated for T-cell recognition are shaded in grey. CVB1_1132-1140_ and CVB3_1135-1143_ differ only by 1 aa and were thus counted as one peptide (total n=16). **G.** CVB nonamer peptides identified in a parallel *in-silico* search for all 6 CVB strains. Homologous β-cell peptides identified by Blast search are indicated with an asterisk. Three peptides that did not confirm as binders *in vitro* were excluded (see <u>Fig. S2</u>). For HLA-eluted peptides, sequence annotations refer to the CVB1/CVB3 strains used to infect β cells. For *in-silico* predicted peptides, annotations indicate sequence identities across serotypes. NA, not applicable or not assigned; ND, not determined.

Three preparations of ECN90 β cells (96-250×10^6^ cells/each) infected with either CVB3 or CVB1 (Fig. 1D) were subsequently analyzed after purification of peptide (p)HLA-I complexes, peptide elution, liquid chromatography – tandem mass spectrometry (LC-MS/MS), and bioinformatics assignment of MS/MS spectra^15^. Altogether, a total of 16 unique CVB peptides were identified, which accounted for a minor fraction (2.9%; range 2.3-3.8%) of the total 8-12 amino acid (aa) peptide display (Fig. 1E-F). The β-cell peptides identified in parallel did not yield any novel hit compared to our previous report^15^ (not shown). A search for putative transpeptidation products generated by the fusion of CVB and β-cell peptide fragments using our previous scripts^15^ did not return any unequivocal assignment.

Only 3/16 peptides (<u>Fig. S1</u>) mapped to the larger structural capsid protein P1 (VP1-VP4); 5/16 mapped to the non-structural protein P3 (3A-D). The smaller non-structural P2 (2A-C) protein comprised most (8/16) of the eluted peptides. The percentage of eluted peptides derived from P2 were significantly more than expected (50% vs. 26%), while few mapped to P1 (19% vs. expected 35-39%; <u>Fig. S2A-B</u>). The eluted peptides were largely conserved across the 6 serotypes, and 5/16 (31%) were eluted from both CVB1- and CVB3-infected β cells.

This list of 16 peptides was complemented: *a)* by searching for length variants mapping to the same region of the eluted peptides^15^ and displaying better HLA-A2/A3-binding score (n=6; indicated in italics in Fig. 1F); *b)* by an *in-silico* prediction of HLA-A2/A3-restricted peptides from representative strains of all 6 CVB serotypes (n=28; Fig. 1G). Of note, CVB1_1356-1364_/CVB3_1359-1367_ (not listed in Fig. 1G) was the only HLA-I-eluted peptide that was also predicted *in silico*, supporting the complementarity of the two strategies.

Minor sequence homologies with β-cell proteins (i.e. IA-2, IGRP, ZnT8) were noted for some CVB peptides (Fig. 1F-G), barring the eluted CVB1_1132-1140_ (<u>K</u>IL<u>PEVKEK</u>) and CVB3_1135-1143_ (<u>K</u>IL<u>PEV</u>R<u>EK</u>), which displayed a high homology with a GAD_272-280_ peptide (<u>K</u>MF<u>PEVKEK</u>; conserved aa underlined) eluted from previous β-cell preparations^15^. Besides displaying a higher homology for GAD_272-280_, CVB1_1132-1140_ proved to be a better epitope in preliminary T-cell analyses (not shown) and was therefore retained for further studies.

We subsequently focused on predicted HLA-A2/A3-restricted peptides and validated their binding *in vitro* (<u>Fig. S2C-D-E</u>), leading to the final selection of 36 candidates for T-cell studies (Fig. 1F-G, second last column): 9 from immunopeptidomics experiments and 27 from *in-silico* predictions; 24 HLA-A2-restricted and 12 HLA-A3-restricted.

Collectively, these results show that CVB-infected β cells downregulate surface HLA-I expression and present few selected HLA-I-bound viral peptides, which significantly overlap between CVB1- and CVB3-infected cells, are largely conserved across serotypes and mostly map to the viral P2 protein.

### Limited recognition of CVB peptides by circulating CD8^+^ T cells of seropositive individuals

We next tested whether the candidate epitopes identified were recognized by circulating CD8^+^ T cells, using frozen/thawed peripheral blood mononuclear cells (PBMCs) from CVB-seropositive healthy adults (<u>Table S1</u>) and combinatorial HLA-A2 or HLA-A3 multimer (MMr) assays^15,17–19^. The reproducibility was satisfactory for measuring both MMr^+^ T-cell frequencies and naïve vs. effector/memory phenotypes (coefficient of variation, CV 21-27%; <u>Fig. S2F-G</u>). As these assays can probe up to 15 MMr specificities per sample, we screened candidate epitopes in 3 panels: two panels for HLA-A2-restricted candidates (n=12+12; Fig. 2A-B) and one panel for HLA-A3-restricted ones (n=12; Fig. 2C). T-cell frequencies segregated into two groups, above or below 1 MMr^+^/10^5^ total CD8^+^ T cells. This cut-off was therefore selected to retain candidate epitopes, as it corresponds to frequencies above the 1-10/10^6^ typically observed for most peptide-reactive naïve CD8^+^ T cells^15,17,18^. Two additional peptides (HLA-A2-restricted CVB1-2-4-5-6_271-279_ and HLA-A3-restricted CVB1_1356-1364_/CVB3_1359-1367_) were retained because their cognate MMr^+^ cells, despite a frequency above this cut-off only in a donor subgroup, displayed a predominant (>60%) effector/memory phenotype (CD45RA^+^CCR7^‒^, CD45RA^‒^CCR7^+^ or CD45RA^‒^CCR7^‒^). Conversely, only some (6/11) peptides returning high cognate T-cell frequencies displayed a similarly predominant effector/memory phenotype. Overall, 7 HLA-A2-restricted candidates (0/4 HLA-eluted and 7/20 *in-silico* predicted) and 6 HLA-A3-restricted candidates (4/6 HLA-eluted and 2/6 *in-silico* predicted) were retained. Given the high homology and cross-reactivity (not shown) between the HLA-A3-restricted CVB1_1356-1364_/CVB3_1359-1367_ (<u>K</u>I<u>NMPMSVK</u>) and CVB2_1361-1369_ (<u>K</u>V<u>NMPMSVK</u>), only the former (HLA-eluted) peptide was further considered.

**Figure 2.**
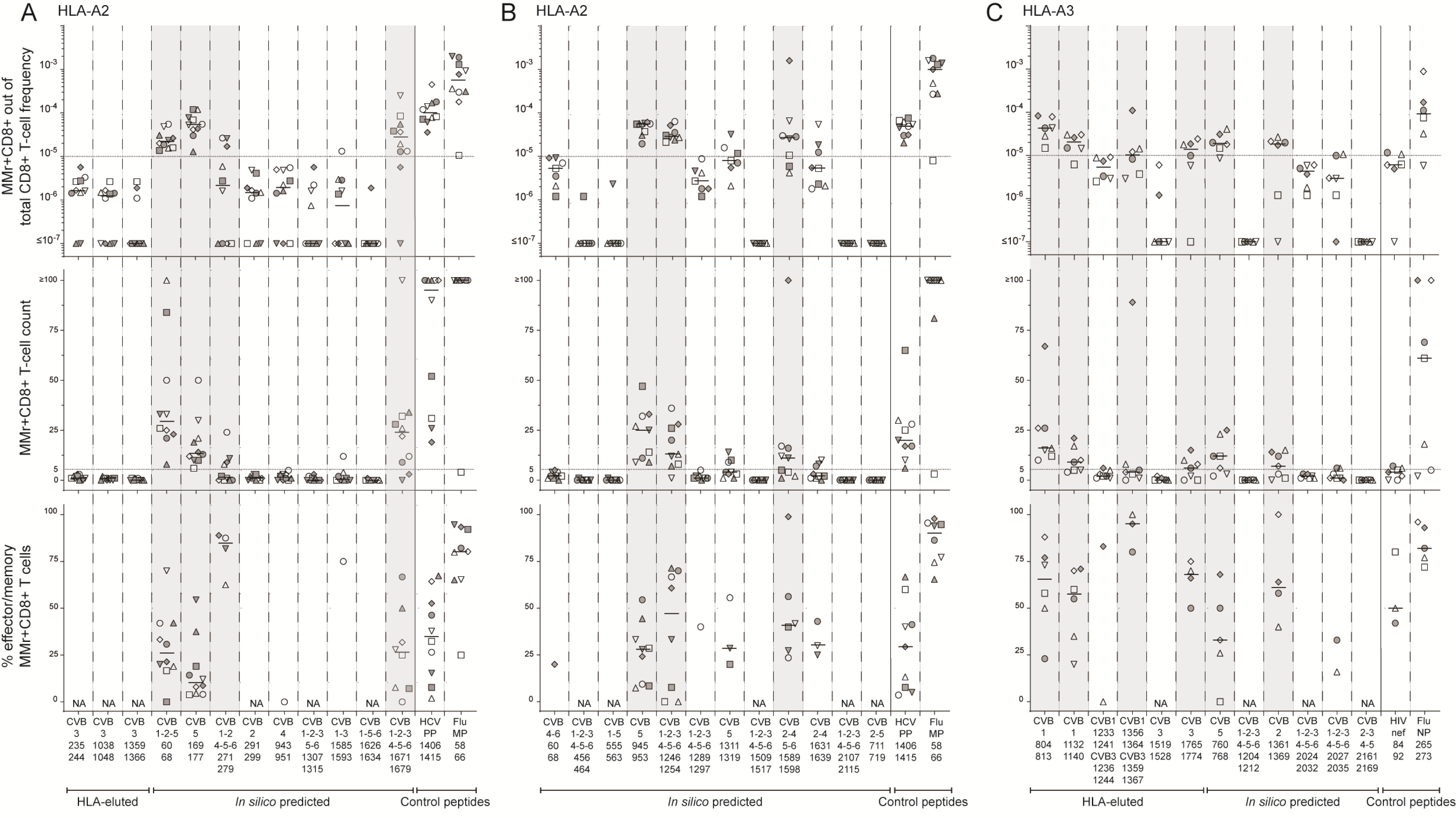
Screening of CVB peptides for recognition by blood CD8^+^ T cells in seropositive healthy adults. **A-C.** HLA-A2-rectricted (A-B) and HLA-A3-restricted candidates (C) were tested with combinatorial MMr assays (see reproducibility in <u>Fig. S2F-G</u>). Each symbol represents a donor (legends in <u>Table S1</u>) and the bars display median values. For each panel, the top graph depicts the frequency of MMr^+^CD8^+^ T cells out of total CD8^+^ T cells, with the horizontal line indicating the 10^−5^ frequency cut-off used as a first validation criterion; the middle graph displays the number of MMr^+^CD8^+^ T cells counted, with the horizontal line indicating the cut-off of 5 cells used to assign an effector/memory phenotype; the bottom graph shows the percent fraction of effector/memory cells (i.e. excluding naïve CD45RA^+^CCR7^+^ cells) among MMr^+^CD8^+^ T cells (for those donors with ≥5 cells counted; NA when not assigned). Peptides validated in this screening phase are highlighted in grey. In each panel, control peptides are derived from viruses eliciting predominantly naïve responses in these unexposed individuals (HLA-A2-restricted HCV PP_1406-1415_ and HLA-A3-restricted HIV nef_84-92_) and from Influenza virus (Flu MP_58-66_ and NP_265-273_ peptides) eliciting predominantly effector/memory responses.

These 12 peptides underwent a second T-cell validation round using more stringent gating and selection criteria. Representative dot plots obtained for these HLA-A2 and HLA-3 MMr panels are shown in <u>Fig. S3</u>. Results are summarized in Fig. 3 and detailed in <u>Table S2</u>. The frequency cut-off was set at 5 MMr^+^/10^6^ total CD8^+^ T cells to account for the more stringent gating criteria applied. Moreover, samples with low total CD8^+^ T-cell counts were excluded to avoid any undersampling bias. Most peptides (5/7 for HLA-A2, 4/5 for HLA-A3; 9/12 in total) were finally validated as relevant targets based on their high cognate T-cell frequency. Of further note, HLA-A3^+^ donors were recruited at two different sites (Paris and Miami) and similar T-cell frequencies were detected in the two subgroups (Fig. 3C).

**Figure 3.**
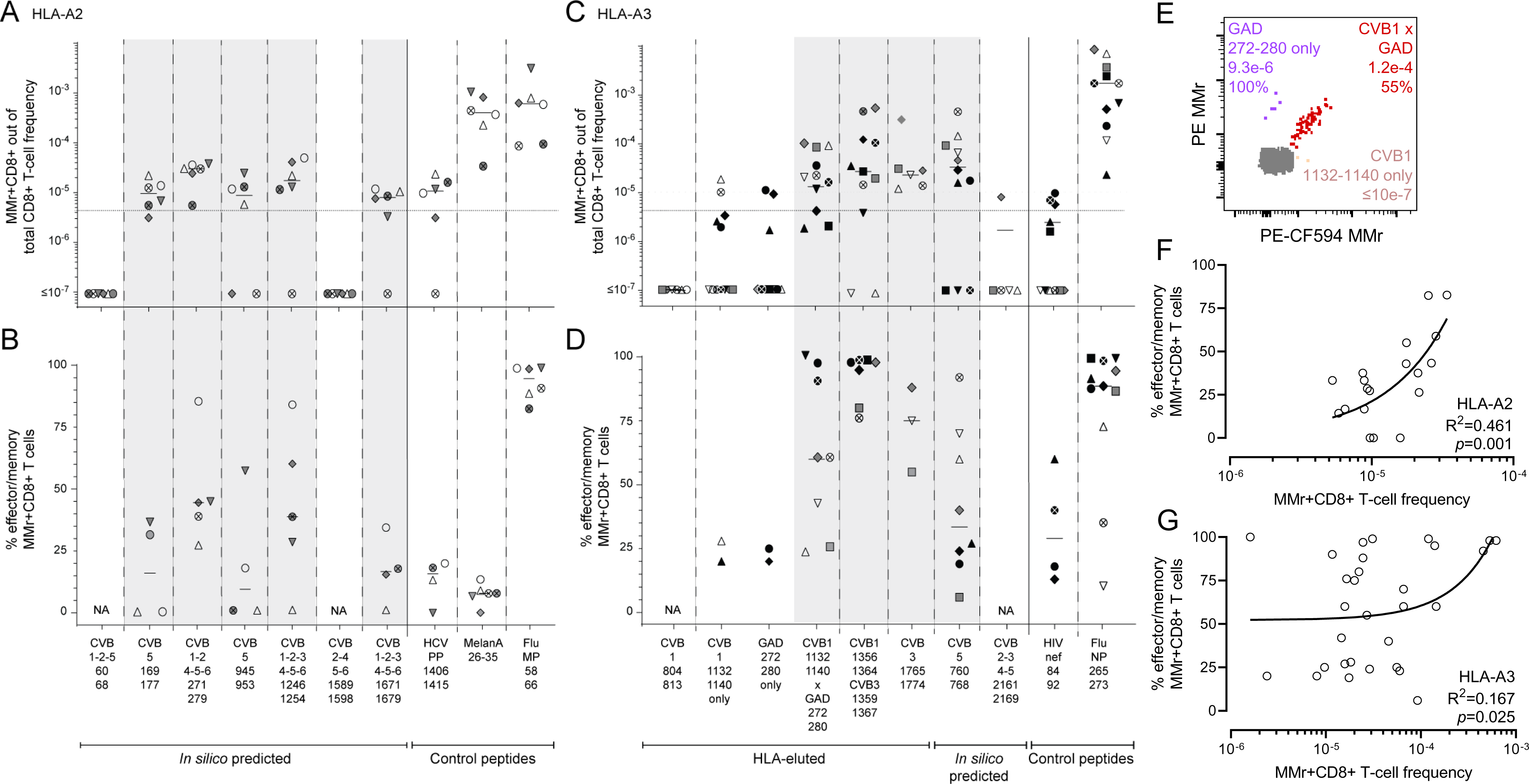
Validation of CVB epitopes as targets of blood CD8^+^ T cells in seropositive healthy adults. **A-D.** The HLA-A2-rectricted and HLA-A3-restricted peptides selected for validation from the screening round (Fig. 2) are shown in panels A-B and C-D, respectively (see <u>Fig. S3</u> for representative dot plots and <u>Table S2</u> for detailed results). In the HLA-A3 panel, the CVB2-3-4-5_2161-2169_ peptide not retained after screening was here included as negative control. Each symbol represents a donor (legends in <u>Table S1</u>), with donors recruited in Paris and Miami depicted in panels C-D as white/gray and black symbols, respectively. Bars indicate median values. Panels A, C depict the frequency of MMr^+^CD8^+^ T cells out of total CD8^+^ T cells, with the horizontal line indicating the 5×10^−6^ frequency cut-off used for validation, with only peptides scoring ≥3 MMr^+^ cells retained for this more stringent analysis. Panels B, D show the percent fraction of effector/memory cells among MMr^+^CD8^+^ T cells. Validated peptides are highlighted in grey. In each panel, control peptides are derived from viruses or self-antigens eliciting predominantly naïve responses (HLA-A2-restricted HCV PP_1406-1415_/MelanA_26-35_ and HLA-A3-restricted HIV nef_84-92_) and from Influenza virus, eliciting predominantly effector/memory responses. For the CVB1_1132-1140_ peptide homologous to GAD_272-280_, frequencies and effector/memory fractions are shown for T cells recognizing either peptide or both. **E.** CD8^+^ T cells cross-reactive to CVB1_1132-1140_ (loaded on PE-CF594/BV711 MMrs) and GAD_272-280_ (loaded on PE/BV711 MMrs) appearing as a triple-stained population labeled by the 4 MMrs (see <u>Fig. S3C-D</u> for gating details). **F-G.** Correlation between the frequency of CVB epitope-reactive MMr^+^CD8^+^ T cells and their percent effector/memory fraction for HLA-A2-(F) and HLA-A3-restricted peptides (G). Each symbol represents an individual epitope specificity in an individual donor.

The HLA-A3-restricted GAD_272-280_ peptide homolog to CVB1_1132-1140_ was further included at this stage to assess any potential cross-reactivity. To this end, we employed our previous assay modification^19^ in which the MMr pair loaded with CVB1_1132-1140_ (PE/CF594- and BV711-labeled) shared the BV711 fluorochrome with the MMr pair loaded with GAD_272-280_ (PE- and BV711-labeled), thus allowing to selectively gate CD8^+^ T cells recognizing only the CVB1 or GAD peptide, or both. Most T cells displayed cross-recognition of the CVB1_1132-1140_/GAD_272-280_ homologous peptides, as shown by co-labeling with the two MMr pairs (Fig. 3E, <u>Fig. S3C-D</u>) and by the higher frequencies of CVB1_1132-1140_/GAD_272-280_ MMr^+^ cells vs. those recognizing either peptide alone (Fig. 3C).

Overall, 3/10 (30%) HLA-eluted peptides were validated as robust T-cell targets. The recognition of only a subset of naturally processed and presented peptides did not reflect an MS selection bias, as the same was observed with *in-silico* predicted peptides (7/26, 27% validated as T-cell targets). Moreover, as observed in T-cell screening experiments, the effector/memory fraction was <50% in most individuals. This pattern contrasted with the higher frequency and near-complete effector/memory phenotype of control Influenza virus-reactive CD8^+^ T cells, while it was similar to that observed for the naïve T-cell epitopes hepatitis C virus (HCV) PP_1406-1415_ and HIV nef_84-92_ (all donors being HCV- and HIV-seronegative). Exceptions were noted for HLA-A3-restricted immunodominant CVB1_1356-1364_/CVB3_1359-1367_-reactive T cells and, to a lesser extent, CVB1_1132-1140_/GAD_272-280_-cross-reactive T cells (Fig. 3C-D). Overall, CVB-reactive CD8^+^ T-cell frequencies loosely correlated with the percent effector/memory fractions observed (Fig. 3F-G), suggesting that higher frequencies may partly reflect prior *in-vivo* expansion.

Collectively, these data show that only a fraction of the few CVB peptides displayed by infected β cells is recognized by circulating CD8^+^ T cells of CVB-seropositive individuals, and that only another subset is targeted by predominantly effector/memory T cells. Moreover, CVB1_1132-1140_-reactive T cells cross-recognize a homologous GAD_272-280_ sequence.

### CVB-reactive CD8^+^ T cells display similar features in the blood of adults and children

The modest CVB-reactive CD8^+^ T-cell responses observed in seropositive healthy adults could reflect the long delay since CVB infections, which are more frequent during childhood. We therefore evaluated these responses in HLA-A2^+^ CVB-seropositive children, with or without T1D, recruited at two different sites (Miami and Paris; <u>Table S1</u>). Given that PBMC numbers from pediatric donors are more limited and vary with age, we preliminarily defined the CV of T-cell frequency measurements when decreasing PBMC numbers are sampled (<u>Fig. S4A-B</u>). The minimal total CD8^+^ T-cell count maintaining a CV<35% was thus defined for each epitope, and used to exclude pediatric samples with insufficient counts, thus limiting undersampling bias. As observed for CVB-seropositive healthy adults, children recruited at the two sites did not display major CD8^+^ T-cell differences and were therefore analyzed altogether (<u>Fig. S4C-D</u>). Overall, the frequency of CVB-reactive CD8^+^ T cells detected in children fell in the same range of that of adult donors (Fig. 3), and was similar between T1D and healthy children, barring a marginally higher frequency of CVB5_945-953_-reactive T cells in the healthy group. The effector/memory fraction of CVB1-2-4-5-6_271-279_-reactive T cells was higher in T1D children. As in adults, Influenza virus-reactive CD8^+^ T cells displayed higher frequencies and effector/memory fractions than CVB-reactive ones. A waning effect from remote CVB exposure is therefore unlikely to explain the limited magnitude of CVB-reactive T-cell responses, as such responses were similar between healthy children and adults.

### CVB-reactive CD8^+^ T cells are also found in spleen and pancreatic lymph nodes and display a PD-1^+^ phenotype

A second reason for the limited presence of circulating CVB-reactive CD8^+^ T cells could be sequestration in lymphoid organs. We therefore searched for these T cells in spleen cell samples from the Network for Pancreatic Organ Donors with Diabetes (nPOD; <u>Table S3</u>, <u>Fig. S5A</u>). Given the small set of samples available, this analysis was not designed to test differences across T1D, aAb^+^ and non-diabetic donors, but rather to assess whether the frequency and/or effector/memory phenotype was different compared to blood. Despite the limited T-cell sampling imposed by the low cell numbers available, cognate T cells were detected in most donors for HLA-A2-restricted CVB1-2-3-4-5-6_1246-1254_ and HLA-A3-restricted CVB1_1356-1364_/CVB3_1359-1367_ (Fig. 4A). Compared to blood, the spleen harbored similar frequencies but higher effector/memory fractions (90-100%; Fig. 4B).

**Figure 4.**
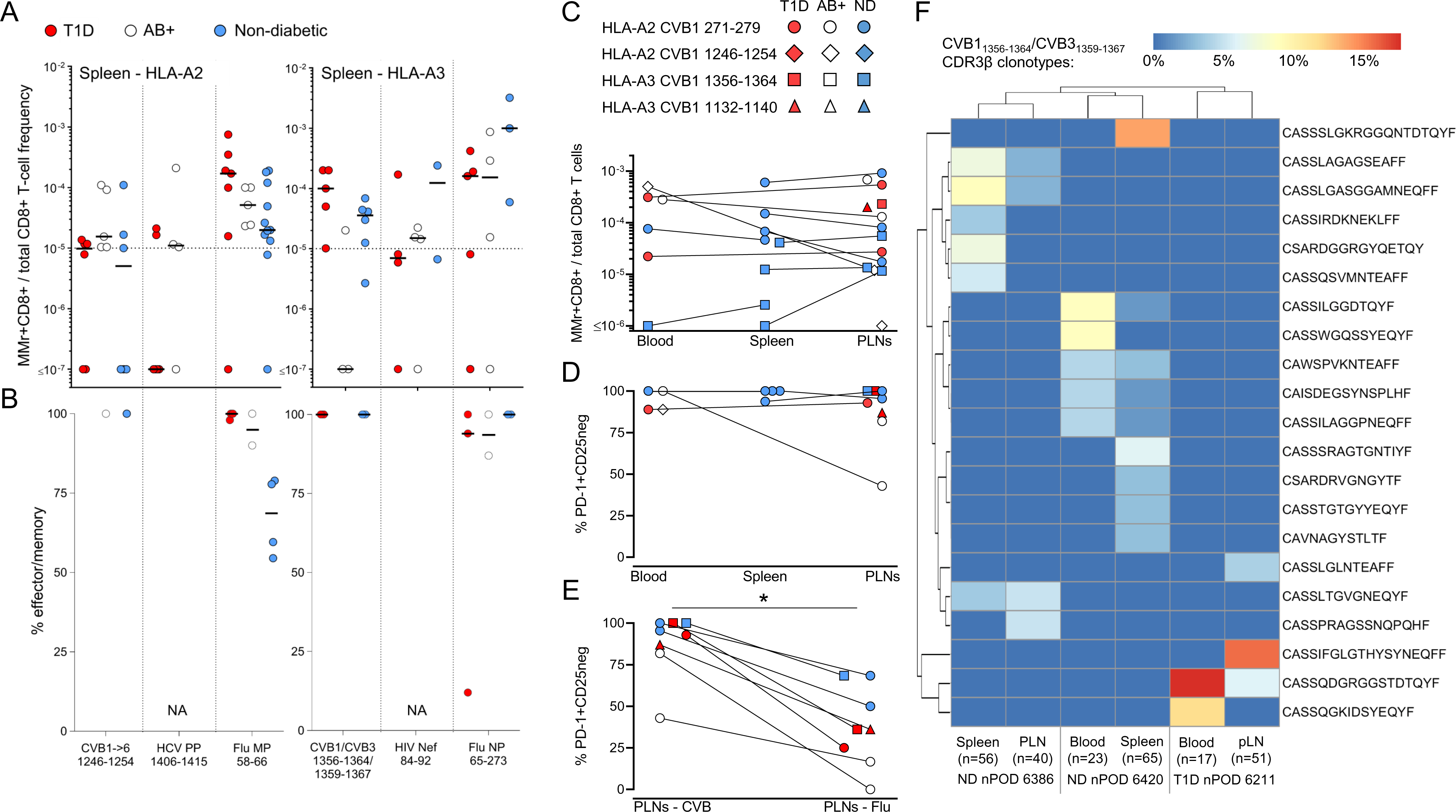
CVB-reactive CD8^+^ T cells are also found in spleen and pancreatic lymph nodes and display a PD-1^+^ phenotype. **A-B.** Detection of CVB epitope-reactive CD8^+^ T cells in splenocytes from nPOD organ donors (HLA-A2^+^, left; and HLA-A3^+^, right; see details in <u>Table</u> <u>S3</u>). Each symbol represents a donor and bars indicate median values. Panel A depicts the frequency of MMr^+^CD8^+^ T cells out of total CD8^+^ T cells, with the horizontal line indicating the 10^−5^ frequency cut-off. A median of 73,320 CD8^+^ T cells were counted (range 9,611-307,697). Only epitopes testing positive are depicted (complete peptide panels listed in <u>Fig. S5A</u>). Panel B displays the percent effector/memory fraction within each MMr^+^CD8^+^ population for those donors/epitopes with ≥5 MMr^+^ cells counted. **C-D.** CVB epitope-reactive CD8^+^ T cells in lymphoid tissues from nPOD cases. CVB MMr^+^CD8^+^ T-cell frequencies (C) and percent PD-1^+^CD25^−^ MMr^+^ cells (D; for donors/epitopes with cell counts ≥5) across available tissues are depicted. **E.** Comparison of PD-1^+^CD25^−^ fractions between CVB and Influenza virus MMr^+^CD8^+^ T cells detected in the same PLN; \**p*=0.014 by Wilcoxon signed rank test. **F.** Expanded TCR CDR3β clonotypes in individual CVB1/CVB3_1356-1364/1359-1367_-reactive CD8^+^ T cells from the same nPOD cases/tissues. Expanded clonotypes were defined as those found in at least 2 MMr^+^ cells in the same tissue from the same donor, and are clustered according to frequency (one columns per tissue/case) and sequence similarity (one row per CDR3β clonotype). Further details are provided in <u>Fig. S5-S6</u>.

For HLA-A3^+^ T1D case #6480 displaying high CVB1_1356-1364_/CVB3_1359-1367_ reactivity (<u>Table S3</u>), available pancreatic lymph node (PLN) cells expanded *in vitro* retrieved CD8^+^ T cells reactive to multiple CVB epitopes (<u>Fig. S5B-C-D-E</u>). We therefore searched for CVB-reactive T cells in the PLNs of nPOD organ donors, and in spleen and PBMC samples when available (<u>Table S3</u>, <u>Fig. S5A</u>; representative staining in <u>Fig. S5F</u>). CD8^+^ T cells reactive to a given CVB epitope displayed similar frequencies across tissues (Fig. 4C). CVB MMr^+^ frequencies were similar to those of Influenza virus-reactive CD8^+^ T cells (<u>Fig. S5G</u>). The vast majority of CVB-reactive T cells displayed an exhausted-like PD-1^+^CD25^−^ phenotype (Fig. 4D), which was instead more limited in control Influenza virus-reactive CD8^+^ T cells (<u>Fig. S5H</u>). Indeed, PD-1 was the most distinctive marker between CVB and Influenza virus MMr^+^ cells across tissues (<u>Fig. S5I</u>). When analyzed within the PLNs of the same donors, CVB-reactive CD8^+^ T cells were also significantly more PD-1^+^CD25^−^ than Influenza virus-reactive ones (Fig. 4E). T-cell receptor (TCR)-sequencing of MMr^+^CD8^+^ T cells reactive to the most immunodominant CVB1_1356-1364_/CVB3_1359-1367_ sorted from paired samples highlighted the presence of expanded CDR3β clonotypes (Fig. 4F). This supports the prior *in-vivo* expansion of these T cells, in agreement with the observed effector/memory phenotype. While some expanded clonotypes were shared across tissues from the same donor, we did not identify public clonotypes (i.e. shared across individuals). Expanded T-cell clonotypes sequenced from PBMCs of CVB-seropositive donors were also exclusively private (<u>Fig. S6A</u>). However, some common CDR3β and CDR3α motifs were noted (<u>Fig. S6B</u>), along with preferential usage of TRBV6 (22%), TRBJ2-1 (22%), TRBJ2-7 (17%), TRBJ2-3 (15%), TRAV8 (19%), TRAV12 (15%) (<u>Fig. S6C</u>) and the pairing of TRBV6 with TRAV8/TRAV12 (13% of total CVB1_1356-1364_/CVB3_1359-1367_ TCRs; not shown).

Collectively, these results show that CVB-reactive CD8^+^ T cells are found in PLNs of T1D, aAb^+^ and non-diabetic donors and harbor expanded private clonotypes and an exhausted-like phenotype.

### CVB-reactive CD8^+^ T cells stained on spleen and PLN tissue sections are more abundant than other viral antigen reactivities in T1D donors

We examined more extensively the frequency and localization of CVB-reactive CD8^+^ T cells on frozen spleen, PLN and pancreas sections from 8 T1D, 2 double-aAb^+^ and 5 non-diabetic donors (<u>Table S4</u>). *In-situ* CD8 staining was combined with MMrs loaded with 2 CVB peptides, CMV/Influenza virus peptides (1/each; memory positive control) or West Nile virus (WNV) peptide (naïve negative control; <u>Fig. S7A</u>). All MMrs were preliminarily validated by *in-situ* staining on purified polyclonal CD8^+^ T cells (<u>Fig. S7B</u>).

In the spleen (Fig. 5A-B-C, Fig. 5J), CVB MMr^+^CD8^+^ T cells were more abundant than their CMV/Influenza virus and WNV counterparts in T1D donors and in the 2 double-aAb^+^ donors available, but not in the non-diabetic group.

**Figure 5.**
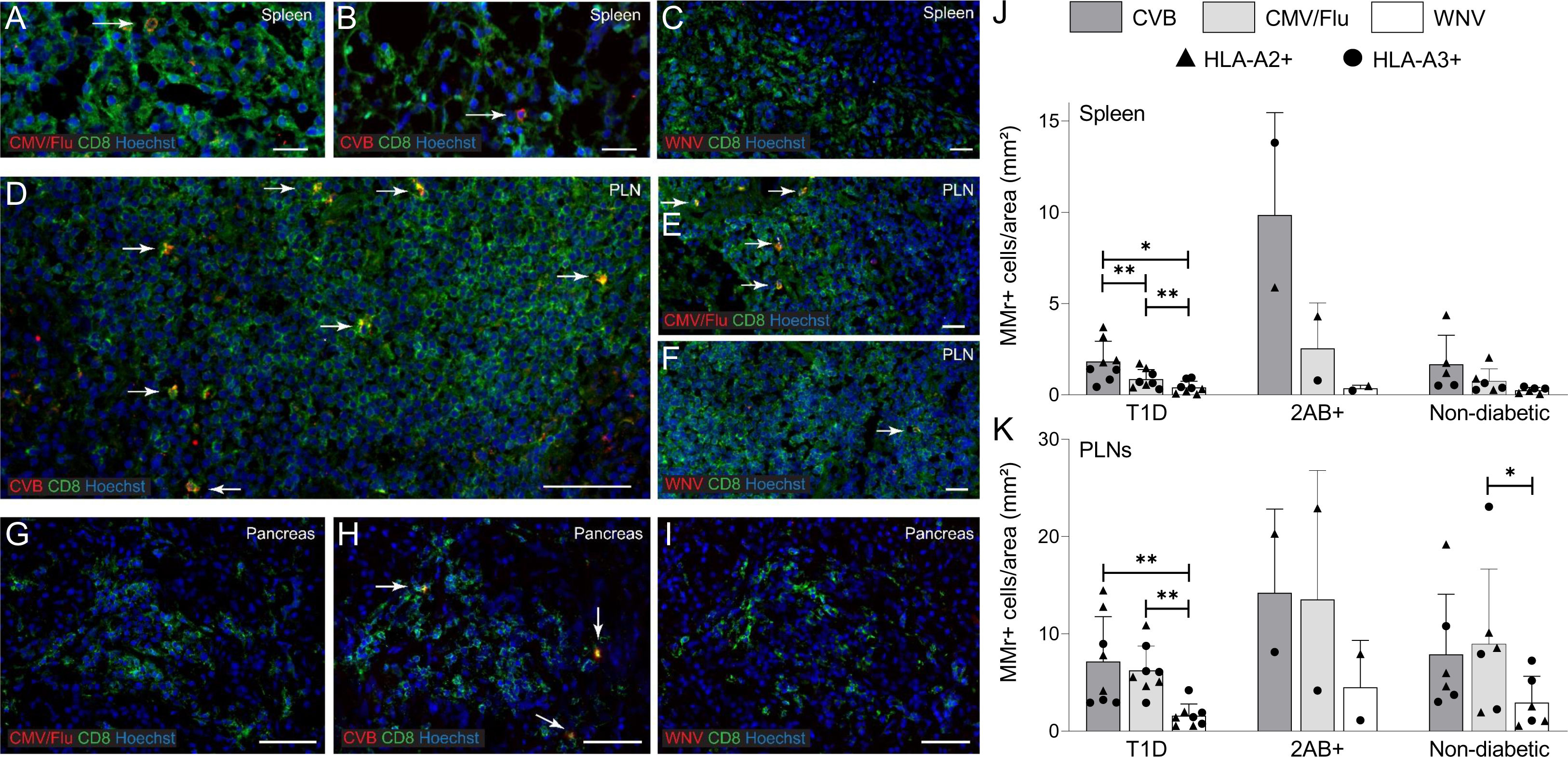
CVB-reactive CD8^+^ T cells stained on spleen and PLN tissue sections are more abundant than other viral antigen reactivities in T1D donors. **A-I.** Representative immunofluorescence images of spleen (A-B-C), PLN (D-E-F) and pancreas (G-H-I) tissue sections (detailed in <u>Table S4</u>) stained with pooled CMV/Flu, pooled CVB or single WNV MMrs (red; see <u>Fig. S7</u>) and CD8 (green). Cell nuclei are stained in blue. Scale bars: 20 μm for spleen, 50 μm for CVB PLN, 20 μm for CMV/Flu PLN, 10 μm for WNV PLN, and 50 μm for pancreas. White arrows indicate MMr^+^CD8^+^ T cells. **J-K.** Bar graphs showing the mean+SD of MMr^+^ cell densities (number of MMr^+^ cells/mm^2^ area) in the spleen (J) and PLNs (K) for CVB, CMV/Flu and WNV specificities in T1D, double-aAb^+^ (2AB+) and non-diabetic HLA-A2^+^ (triangles) and HLA-A3^+^ (circles) donors. \*\**p*=0.008 and \**p*=0.05 by Wilcoxon signed rank test.

In PLNs (Fig. 5D-E-F, Fig. 5K), densities were overall higher than in the matched spleens. CVB-reactive T cells were more abundant than WNV-reactive ones in T1D donors and in the 2 double-aAb^+^ donors available, but not in the non-diabetic group. Control CMV/Flu MMr^+^CD8^+^ T cells were instead more abundant than for WNV in all groups. The overall concordance between CVB MMr^+^ frequencies measured by flow cytometry and CVB MMr^+^ densities quantified by tissue immunofluorescence was weak at low frequencies/densities, but more consistent at higher values (i.e. in double-aAb^+^ donors) (<u>Fig. S7C</u>).

Pancreas sections from the same donors displayed low numbers of islet-infiltrating CD8^+^ T cells in the T1D group, thus impeding an accurate quantification. However, very few or no islet-infiltrating cells were reactive to viral epitopes (Fig. 5G, H, I). Accordingly, CVB-reactive CD8^+^ T cells were not found in the pancreas when assessed by CVB peptide recall assays on carrier T cells transduced with TCRs sequenced from islet-infiltrating CD8^+^ T cells of T1D donors^20^ (not shown).

Collectively, these results show that, while nearly absent in the pancreas, CVB-reactive CD8^+^ T cells are more abundant than other viral antigen reactivities in the spleen and PLNs of T1D donors compared to non-diabetic donors.

### CVB -infected β cells make filopodia and their subsequent death is primarily viral-mediated

We next investigated the mechanisms of β-cell death upon CVB infection. We first assessed the kinetics of CVB infection and β-cell death in the absence of T cells using prolonged real-time imaging and a CVB3-eGFP strain (Fig. 6A). Both infection and death kinetics were accelerated with increasing MOIs. Interestingly, at an intermediate 100 MOI, a first infection plateau was observed at ∼12 h, followed by a higher one at ∼20 h, possibly reflecting the spreading of new virions from a first pool of infected β cells to a new pool. At all MOIs, β-cell death was complete ∼20 h after the infection peak. We also observed that intact β cells acquired CVB upon contact with filopodia protruding from infected cells, subsequently making filopodia themselves and infecting additional cells before dying (Fig. 6B, <u>Movie S1</u>). Accordingly, the eccentricity (i.e. deviation from circularity to quantify cell elongation) of infected β cells varied over time, while it remained stable in non-infected cells (Fig. 6C). VP1^+^ β cells from pancreas tissue sections of T1D donors displayed a similar morphology (Fig. 6D) and higher cell area, perimeter and diameter compared to VP1^−^ ones (Fig. 6E-F-G).

**Figure 6.**
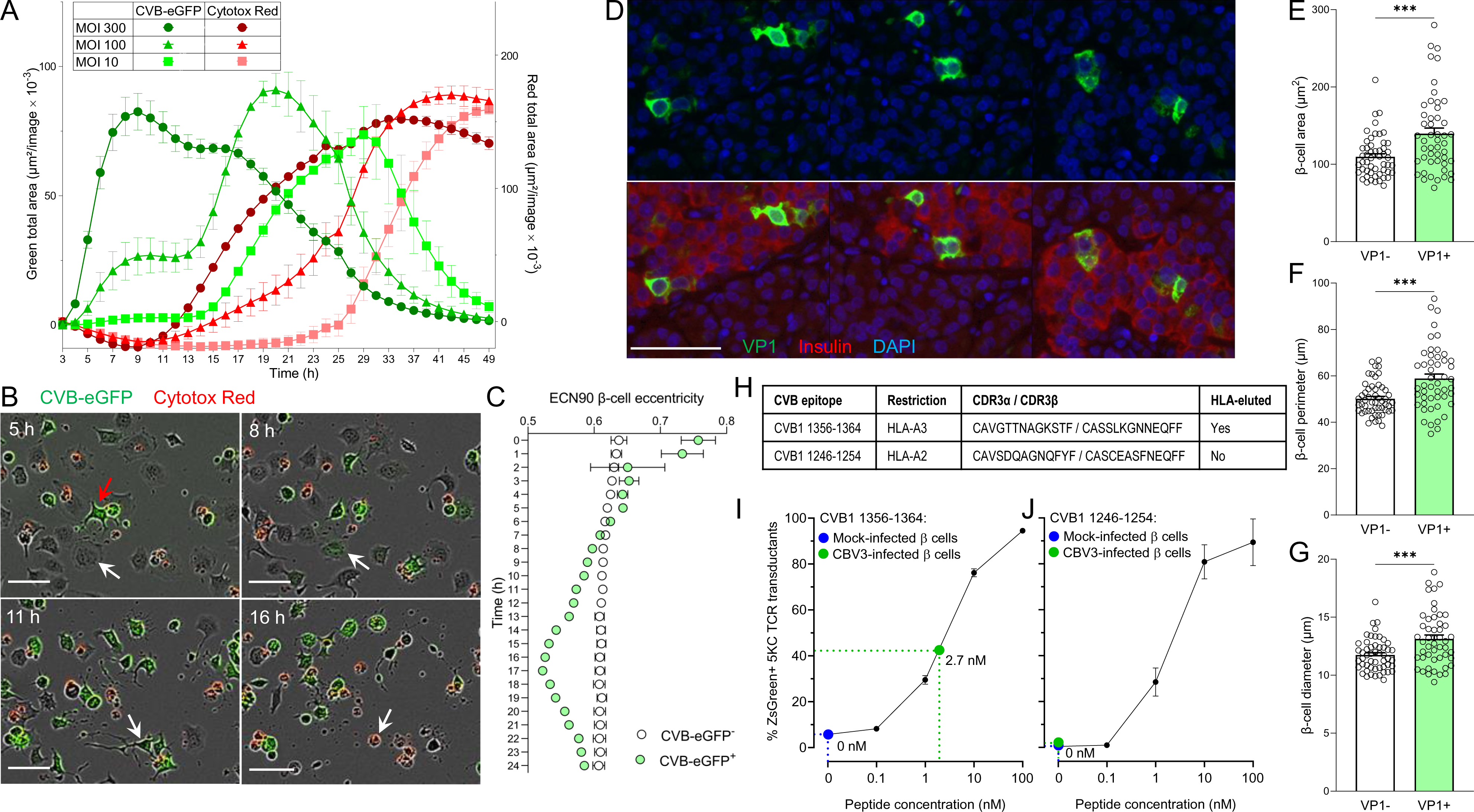
Kinetics of CVB infection and β-cell death and presentation of CVB peptides. **A.** Kinetics of infection and death measured by real-time imaging in ECN90 β cells infected with CVB3-eGFP at the indicated MOI and stained with Cytotox Red to visualize dead cells. Infection curves (green total area, in green) and death curves (red total area, in red) are plotted on the left and right y-axis, respectively. Each data point represents the mean±SD of at least duplicate measurements from a representative experiment performed in duplicate. **B.** Representative images from <u>Movie S1</u> showing an intact β cell at 5 h (white arrow) in contact with the filopodia of an infected eGFP^+^ cell (red arrow), turning eGFP^+^ at 8 h and subsequently emanating filopodia at 11 h before dying (Cytotox Red^+^) at 16 h. Scale bar: 30 μm. **C.** Eccentricity of infected vs. non-infected ECN90 β cells (300 MOI). Each point represents the mean±SD of 8 images, *p*<0.01 at all time points (barring 2 h) by unpaired Student’s t test. **D.** Representative immunofluorescence images from pancreas tissue sections of T1D donor 6243 stained for VP1 (green) and DAPI (blue), alone (top) or after merging with insulin staining (red, bottom). Scale bar: 50 µm. **E-F-G.** Comparison of VP1^−^ vs. VP1^+^ β-cell area (E), perimeter (F) and diameter (G) in pancreas tissue sections stained as above. Each point represents a β cell from sections obtained from 3 T1D donors; bars represent mean+SEM values. \*\*\**p*≤0.0004 by unpaired Student’s t test. **H.** CVB-reactive TCRs sequenced and re-expressed in carrier cells. **I.** CVB1_1356-1364_-reactive TCR activation in NFAT-driven ZsGreen fluorescent reporter, TCR-transduced 5KC cells upon an 18-h co-culture with CVB3-infected (green; 300 MOI) vs. mock-infected ECN90 β cells (blue), plotted as mean ± 95% confidence interval of triplicate wells from a representative experiment performed in duplicate. A parallel dose-response curve with increasing peptide concentrations pulsed on non-infected β cells (black) estimated the concentration of peptide recognized on CVB-infected β cells to 2.7 nM. **J.** Negative control TCR recognizing the CVB1_1246-1254_ peptide (not eluted from β cells) transduced into the same reporter 5KC cells and co-cultured as above, depicted as in panel I.

To study how CVB-reactive CD8^+^ T cells contribute to β-cell death, an immunodominant TCR clonotype recognizing HLA-A3-restricted CVB1_1356-1364_/CVB3_1359-1367_ (Fig. 6H) was re-expressed into carrier cells. TCR-transduced 5KC T cells exposed to CVB3-infected β cells upregulated expression of a fluorescent NFAT-induced ZsGreen reporter^21^, indicating display of the cognate peptide (Fig. 6I). A parallel dose-response standard curve using peptide-pulsed non-infected β cells estimated the concentration of CVB peptide naturally processed and presented by infected β cells to 2.7 nM. Thus, the natural CVB antigen presentation is sizable and sufficient to stimulate CVB-reactive T cells. A TCR recognizing HLA-A2-restricted CVB1-2-3-4-5-6_1246-1254_ (not eluted from β cells; Fig. 6H) did not elicit T-cell activation upon contact with CVB-infected β cells in the absence of prior peptide pulsing (Fig. 6J).

Cytotoxic primary CD8^+^ T cells were then transduced with the same CVB1_1356-1364_/CVB3_1359-1367_ TCR and exposed to infected ECN90 β cells. Real-time imaging revealed that CVB-induced β-cell death (i.e. in the absence of T cells) had distinctive morphological features compared to death induced by T cells (i.e. non-infected β cells pulsed with CVB peptide). While CVB-infected β cells died in a discrete, single-cell pattern (Fig. 7A, <u>Movie S2</u>), those attacked by T cells formed clusters (Fig. 7B, <u>Movie S3</u>). These distinct patterns allowed us to create single-cell and cluster-cell analysis masks to differentially quantify CVB- and T-cell-induced death, respectively (<u>Movie</u> <u>S4A-B</u>). Using this strategy, we compared the percentage of CVB-killed vs. T-cell-killed β cells over 48 h, using increasing MOIs and T:β cell ratios (Fig. 7C). As expected, both types of cell death increased with time. While CVB killed β cells earlier, T cells started to do so only after ∼24 h. Starting from this time point, we therefore studied the relative contribution of CVB- and T-cell-induced death (Fig. 7D; <u>Fig. S8A</u>). CVB-mediated killing increased with higher MOIs in the absence of T cells, but also in their presence for T:β-cell ratios up to 1:1. T-cell-mediated killing became predominant only at high T:β-cell ratios of 2:1; however, this also reflected a decreased CVB-mediated killing. Since this decrease started already before 24 h (Fig. 7C) and was equivalent when using non-transduced, non-CVB-reactive CD8^+^ T cells (<u>Fig. S8B-C-D</u>), it likely reflects reduced spreading of CVB from infected to intact β cells in the presence of high T-cell numbers, independent of their cytotoxicity. There was no measurable infection of T cells. Collectively, these results show that most death of infected β cells is primarily induced by CVB. T-cell-mediated cytotoxicity becomes predominant only at high numbers, which also reflects a steric limitation of CVB spreading by T cells, independently of their antigen reactivity.

**Figure 7.**
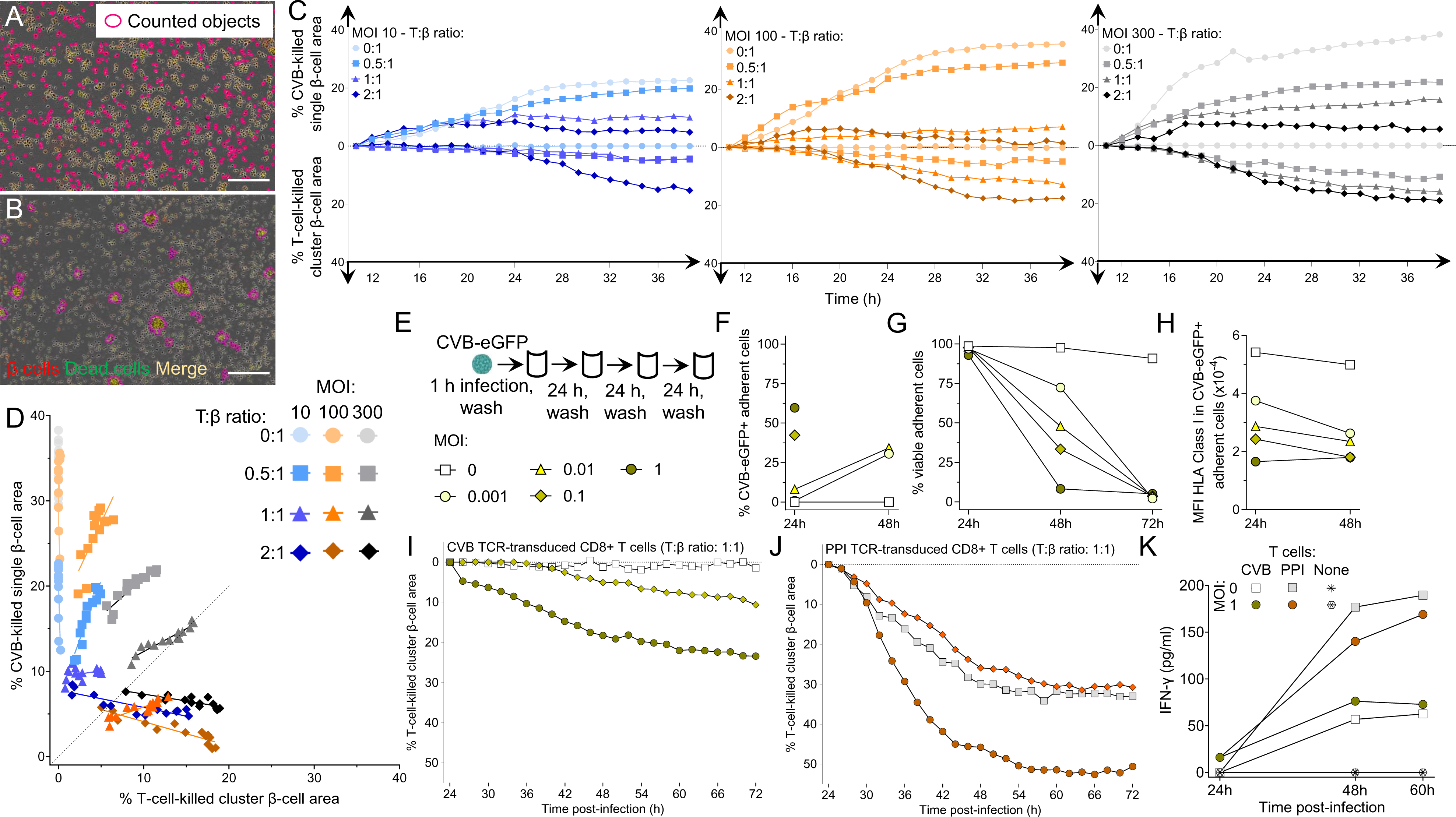
Infected β cells are more efficiently killed by CVB than by CVB-reactive T cells. **A-B.** Representative real-time images from <u>Movie S4A-B</u> (taken at 300 MOI, 1:1 T:β-cell ratio) showing (A) CVB-killed β cells detected by single-cell analysis mask and (B) β cells killed by CVB TCR-transduced CD8^+^ T cells, detected by cluster analysis mask. β cells and dead cells were labeled with CellTracker Red and Cytotox Green dye, respectively; dead β cells are therefore labeled in yellow. Scale bar: 30 μm. CVB-killed single β cells counted in the absence of T cells and T-cell-killed cluster β cells counted in the absence of CVB after peptide pulsing are shown in <u>Movies S2-S3</u>, respectively. **C.** ECN90 β cells were stained as above and infected with CVB at the indicated MOIs. Curves display the percent CVB-killed single β-cell area (top) and the percent T-cell-killed cluster β-cell area (bottom) at the indicated T:β-cell ratios over a 48-h real-time imaging acquisition. β-cell areas are expressed as median percent of total β-cell area from at least duplicate wells, normalized to the 8-h time point. A representative experiment out of 2 performed is displayed. **D.** Correlation analysis between the percent CVB-killed single β-cell area and the percent T-cell-killed cluster β-cell area from the same experiment depicted in (C) during the last 24 h of acquisition. Further details are provided in <u>Fig. S8</u>. **E.** Schematic of the low-grade infection protocol. ECN90 β cells were incubated with CVB3-eGFP for 1 h, followed by washing and culture for 72 h, with washes to remove free virions, cell sampling for flow cytometry and medium replenishment every 24 h. **F.** Percent CVB3-eGFP^+^ adherent ECN90 β cells detected at 24 and 48 h at different MOIs. **G-H.** Percent viable adherent cells (G) and HLA-I MFI (H) in CVB3-eGFP^+^ adherent cells from the same experiment. Each point represents the mean of duplicate measurements from a representative experiment. For panels F and H, some time points are not depicted due to the low number of remaining viable cells. **I-J.** Percent T-cell-killed cluster β-cell area following infection for 1 h at the indicated MOIs, washing, 24-h culture and real-time imaging after addition of CVB1_1356-1364_ (I) or PPI_15-24_ (J) TCR-transduced CD8^+^ T cells at a 1:1 T:β cell ratio. β-cell areas are expressed as median percent of total β-cell area from at least duplicate wells of a representative experiment, normalized at the 24-h time point. Symbol legends are the same as in panel E (depicted in different colors for panel J). **K.** IFN-γ secretion at the indicated MOIs and T-cell/β-cell co-culture conditions. Each point represents the mean of duplicate measurements from the experiment depicted in panels I-J.

To investigate whether low-grade CVB infection leads to similar outcomes, ECN90 β cells were infected with CVB-eGFP at low MOIs (0.001, 0.01, 0.1 and 1; Fig. 7E) and analyzed for the kinetics of infection, β-cell death and HLA-I modulation over 72 h. Both infection (Fig. 7F) and death (Fig. 7G) increased with time in a MOI-dependent manner, with complete β-cell death at all MOIs by 72 h. HLA-I down-regulation was also observed at these low MOIs (Fig. 7H). We then infected β cells at 0.1 and 1 MOIs and, after 24 h (when viability was still intact), we added CD8^+^ T cells transduced with either the previous CVB1_1356-1364_/CVB3_1359-1367_ TCR or a β-cell-reactive HLA-A2-restricted preproinsulin (PPI)_15-24_ TCR^22^ for real-time imaging. Also under this low-grade infection conditions, the killing mediated by CVB-reactive T cells was limited, plateauing at 23% (Fig. 7I), while PPI-reactive T-cell-mediated killing was more extensive (Fig. 7J), thus excluding a general β-cell resistance to cytotoxicity induced by CVB, e.g. by HLA-I downregulation. Moreover, the higher MOI 1 enhanced cytotoxicity of both CVB- and PPI-reactive T cells, but not their IFN-γ secretion (Fig. 7K), suggesting that this enhanced cytotoxicity reflects increased β-cell vulnerability rather than increased T-cell activation. IFN-α2a, IFN-β and IFN-λ remained undetectable throughout the culture period (not shown).

Collectively, these *in-vitro* results indicate that, upon infection, β cells are mostly killed directly by CVB, with a limited contribution of CVB-reactive CD8^+^ T cells. CVB infection can however enhance the β-cell death mediated by autoimmune PPI-reactive CD8^+^ T cells.

## Discussion

Despite lack of evidence for a causal relationship, CVB infections are gaining credit as candidate environmental triggers for T1D based on their temporal correlation with aAb seroconversion^6^ and on detection of pancreatic hallmarks of infection predominantly in T1D patients^8–10^. These associations have prompted the development of a multivalent CVB1-6 vaccine^11,23^, which formed the basis for a phase I trial in healthy adults of a PRV-101 vaccine covering the most prevalent CVB1-5 serotypes (NCT04690426). It is therefore relevant to address the knowledge gaps on CVB-induced immunity. While Ab responses have been extensively studied, T-cell responses are poorly characterized^1^. Such information would contribute to understanding the mechanisms of CVB-induced β-cell damage and inform the development of vaccine trials. We focused on CD8^+^ T cells, which are key players in both anti-viral responses and in the autoimmune attack against β cells. Two different, non-mutually exclusive scenarios could explain how CVB triggers T1D^1^. First, weak anti-CVB immune responses could favor viral spreading and persistence, resulting in an enhanced, primary CVB-mediated killing of infected β cells. Second, an opposite scenario of strong anti-viral immune responses could lead to indirect, secondary killing of infected β cells by viral-reactive T cells. In all cases, the final outcome would be the release of self-antigens in an inflammatory environment that may initiate/amplify autoimmunity, but the implications for vaccination strategies are different. While CVB vaccination may be beneficial to enhance weak immune responses and limit viral spreading, the second scenario may invite caution, as boosting anti-viral T-cell responses could trigger immune-mediated destruction of infected β cells.

The current study fills some of these gaps with several findings. First, an efficient immune escape mechanism mounted by CVB is down-regulation of surface HLA-I on infected β cells, as previously reported^24,25^, which leads to the presentation of very few, selected CVB peptides. Most of them mapped to non-structural proteins P2-P3, despite the fact that they lie downstream of structural P1 (VP1-VP4) in the viral polyprotein and are thus translated last. Non-structural proteins may be better substrates for epitope display because they are not encapsulated in newly-formed virions, thus making them more readily accessible for antigen processing and presentation. Second, CVB transfer through filopodia, previously described in other cells^26^, may provide another immune escape mechanism by shielding virions from immune recognition and Ab-mediated neutralization.

Third, the overall strength of anti-CVB CD8^+^ T-cell responses is limited in terms of number of epitopes recognized out of the few presented by β cells. While effector/memory responses were documented by surface phenotype and expanded TCR clonotypes, they were limited for most epitopes and donors, as supported by the low T-cell frequencies observed, by a substantial naïve fraction, and by an exhausted-like PD-1^+^ phenotype, as observed in other persistent viral infections^27,28^. This poor memory does not seem to reflect a long delay since CVB exposure, as T-cell frequencies and effector/memory fractions in adults were similar to those in children, who are more frequently infected by CVBs. Moreover, poor memory generation seems universal, as similar CVB-reactive T-cell frequencies and effector/memory fractions were observed in T1D and healthy children. This is not unexpected, in light of the late stage 3 disease of these T1D patients. Studies on longitudinal samples from earlier T1D stages are needed to assess whether anti-CVB T-cell memory is further diminished in children later developing islet aAbs, as observed in serological studies suggesting poor anti-CVB Ab responses^12,29^. This could favor the prolonged pattern of CVB infections associated with islet aAb seroconversion^6^. Footprints of CVB presence are also observed in T1D pancreata^8–10^. On the other hand, the density of CVB-reactive CD8^+^ T cells was higher than that of other viral specificities in the spleen and PLN tissues of T1D donors, suggesting that, in the disease setting, these responses may be more efficiently elicited in secondary lymphoid organs, possibly in the context of higher CVB loads gaining access to the pancreas.

Collectively, these observations provide some rationale to induce immune memory through vaccination. Although CVB-reactive CD8^+^ T cells were capable of killing infected β cells, their role was predominant only at the highest T:β cell ratios and was favored by a concomitant steric inhibition of CVB transfer. Killing was otherwise mainly CVB-mediated, even at low MOIs, suggesting that autoimmune triggering predominantly relies on the direct, primary cytotoxic effect of the virus releasing islet antigens. In line with these observations, CVB-reactive CD8^+^ T cells were near-absent in islet infiltrates. These results suggest that vaccination is unlikely to enhance β-cell destruction, as ruled out in rhesus macaques^11^. In addition, the multivalent PRV-101 CVB vaccine is produced with chemically inactivated mature virions that do not comprise the non-structural peptides preferentially presented by β cells, including the CVB1_1132-1140_/CVB3_1135-1143_ epitope cross-reactive with the endogenous β-cell peptide GAD_272-280_. This peptide maps to a larger GAD region, whose CVB cross-reactivity initially reported^30^ was subsequently dismissed^31,32^. We now show that CVB/GAD cross-recognition is at play for CD8^+^ T cells, with the large majority of them reactive to both epitopes. To better understand the pathogenic mechanisms of CVB infection, it will be important to test whether these cross-reactive T cells can lyse non-infected β cells. However, the opposite possibility that these CVB/GAD cross-reactive responses may actually provide better anti-viral protection should not be dismissed^33^. Indeed, anti-GAD aAbs are more prevalent at older ages^34^ and have been suggested to be protective, as children with multiple aAbs, including anti-GAD, have a lower risk of clinical progression than those without anti-GAD^33^.

By combining immunopeptidomics and *in-silico* predictions, we identified several other epitopes that, although not presented by β cells, were recognized by CD8^+^ T cells. Overall, we identified two epitopes for each of the HLA-A2 and HLA-A3 restrictions studied that displayed high immunodominance and immunoprevalence: HLA-A2-restricted CVB1_271-279_, CVB1_1246-1254_ and HLA-A3-restricted CVB1_1132-1140_, CVB1_1356-1364_. A confounding effect of polio vaccination is unlikely, in light of their limited homology with the vaccine poliovirus serotypes (<u>Fig. S9A</u>); one reason for their preeminence may instead be their conservation across CVB serotypes. Importantly, a larger panel of epitopes (<u>Fig. S9B</u>) mapping to structural P1 (VP1-4) (e.g. HLA-A2-restricted CVB5_169-177_, CVB1_271-279_; HLA-A3-restricted CVB5_760-768_) and non-structural P2-P3 (e.g. HLA-A2-restricted CVB1_1246-1264_, CVB1_1671-1679_; HLA-A3-restricted CVB1_1132-1140_, CVB1_1356-1364_, CVB3_1765-1774_) could be used to differentially monitor T-cell responses to natural infection (where both structural and non-structural epitopes should be targeted) vs. those elicited by vaccination (with only structural epitopes recognized), analogous to what is routinely done for hepatitis B virus (by testing anti-HBsAg vs. anti-HBcAg Abs) and SARS-CoV-2 (by testing anti-spike vs. anti-nucleoprotein Abs). This distinction cannot be made by CVB serology.

The surface HLA-I downregulation induced in CVB-infected β cells decreased the presentation not only of CVB peptides, but also of endogenous ones. Indeed, only 152-206 8-12mer human peptides were retrieved from each biological replicate (96-250×10^6^ cells) as compared to the 3,544 peptides retrieved using equivalent or lower total cell numbers (100×10^6^) of untreated β cells in our previous work^15^. However, besides causing β-cell lysis and self-antigen release, CVB infection triggers a type I IFN response^1^, which may create an immunogenic environment and induce the secondary HLA-I hyper-expression co-localized with viral VP1 protein in islets of T1D patients^9^ and persisting years after clinical diagnosis^35^. This HLA-I hyperexpression leads to an increased peptide presentation^15^, thus enhancing β-cell vulnerability to autoimmune T cells. While we observed an enhanced cytotoxicity of PPI-reactive CD8^+^ T cells against CVB-infected β cells, this enhancement was not paralleled by an increased IFN-γ secretion, nor by any detectable IFN-α2a, IFN-β or IFN-λ, suggesting that other potentiation mechanisms may be at play. These *in-vitro* results should however be interpreted with caution, as IFN responses may originate not only from T cells (IFN-γ) and β cells (mainly IFN-β), but also from plasmacytoid dendritic cells (mainly IFN-α) responding to DNA from dying β cells^36^ and, possibly, viral RNA. Multivariate *in-vitro* systems, e.g. using pancreas tissue slices, are needed to more accurately model these interactions. However, IFNs may be insufficient at mitigating CVB evasion from HLA-I presentation, and rather favor their persistence hidden inside β cells. Indeed, the group of L. Whitton^37^ used *in-vivo* mouse models to elegantly show that a recombinant CVB3 encoding highly immunogenic lymphocytic choriomeningitis virus (LCMV) epitopes barely activates LCMV TCR-transgenic T cells, suggesting near-complete evasion from MHC Class I presentation even when IFN responses are operational.

This study carries limitations. First, only the peptides naturally processed and presented by β cells were considered. While this allowed us to directly address the possibility that vaccination may trigger CD8^+^ T-cell cytotoxicity against β cells, the peptide display offered for T-cell activation by other cells, including antigen-presenting cells, may be broader, as suggested by the identification of additional epitopes by *in-silico* predictions. Notably, it will be important to define the peptides displayed by infected enterocytes at the viral entry site, and by dendritic cells and macrophages that may take up antigens locally and in draining lymph nodes. Second, the contribution of CD4^+^ T-cell responses to CVB immunity remains to be explored. Third, it is possible that the interplay between CVB, β cells and T cells described herein may change if the primary infection evolves toward a chronic, non-replicative persistence in β cells, e.g. by naturally occurring 5’-terminal deletions of the CVB genome^38^.

In conclusion, the identification of the CVB epitopes targeted by CD8^+^ T cells allowed us to define the role of CVB infection in mediating primary β-cell lysis while inducing limited anti-viral CD8^+^ T-cell responses, which mitigates the risk that CVB vaccination may precipitate the destruction of infected β cells. The poor CD8^+^ T-cell memory against CVB lends rationale for inducing such memory through vaccination.

## Acknowledgments.

We thank the Cochin Institute CyBio Flow Cytometry Core Facility for assistance with cell analysis and sorting. We gratefully acknowledge Dr. Vesa Hytönen (University of Tampere, Finland) for the gift of the anti-VP1 mAb 3A6; Dr. Lindsay Whitton (The Scripps Research Institute, La Jolla, CA, USA) for the gift of the CVB3-eGFP strain; Dr. Sami Oikarinen and Dr. Minna Hankaniemi (University of Tampere, Finland) for CVB sequencing and dynamic light scattering quality controls; and Dr. Mark Peakman (King’s College, London) for providing the 1E6 TCR sequence.

## Author Contributions

Conceptualization: F.V., A.C., S.T., I.S., S.Y., A.P., H.H., T.R.C., M.F.T., R.M.; Methodology: F.V., A.C., D.K., Z.Z., P.A., S.T., I.S., O.B.M., J.P.H., B.B., G.A., C.H., J.K., S.C.K., M.N., S.J.R., J.V., Y.V., R.S., S.B., S.Y., A.P., H.H., T.R.C., M.F.T., R.M.; Investigation: F.V., A.C., D.K., Z.Z., P.A., S.T., I.S., O.B.M., J.P.H., B.B., G.A., C.H., J.K., S.C.K., M.N., S.J.R., J.V., Y.V., J.L., M.S., Z.M., E.B., N.L., J.S., C.B., P.G.; Visualization: F.V., A.C., D.K., Z.Z., P.A., O.B.M., J.P.H., C.H., J.K., S.C.K., S.J.R., T.R.C., R.M.; Supervision: J.V., S.Y., A.P., T.R.C., M.F.T., R.M.; Funding Acquisition: A.P., H.H., T.R.C., M.F.T., R.M.; Manuscript Writing: F.V., A.C., T.R.C., M.F.T., R.M.

## STAR Methods

### Preparation and titration of CVB1 and CVB3 stocks

CVB1 (ATCC VR-28 strain), CVB3 (Nancy strain) and CVB3-eGFP^39^ were grown in HeLa cells. Supernatants and cells were collected after overnight culture, subjected to 3 freeze/thaw cycles, centrifuged at 4,000 rpm and further purified by ultracentrifugation. Next-generation sequencing was used to confirm serotypes and identify sequence variants (quasi-species). Virus stocks were titrated on HeLa cells.

### *In-vitro* infection

The ECN90 β-cell line (HLA-A*02:01/03:01, HLA-B*40:01/49:01, HLA-C*03:04/07:01)^14,17^ was cultured in DMEM/F12 Advanced medium (ThermoFisher) supplemented with 2% bovine serum albumin (BSA) fraction V, fatty-acid-free (Roche), 50 µM β-mercaptoethanol (Sigma), 10 mM nicotinamide (Merck), 1.7 ng/mL sodium selenite (Sigma), 100 U/mL Penicillin-Streptomycin (Gibco), 1% glutaMAX (Gibco). Cells were seeded at 16.5×10^6^ cells in 150-cm² culture flasks (TPP) coated with 0.25% fibronectin from human plasma (Sigma) and 1% extracellular matrix from Engelbreth-Holm-Swarm murine sarcoma (Sigma) and cultured at 37°C in 5% CO_2_ for 16-24 h. They were infected with 300 MOI of CVB1 or CVB3 for 1 h in BSA-free supplemented medium. After gentle washing, the culture was pursued for 6 h, followed by trypsinization, washing with phosphate-buffered saline (PBS) and processing for flow cytometry or freezing at –80°C as dry pellets for peptidomics experiments.

Infection was verified on cells stained with Live/Dead Violet (ThermoFisher) and monoclonal Abs (mAbs) to HLA-A,B,C (RRID:AB_2917755), VP1 (clone 3A6)^40^, dsRNA (RRID:AB_2651015) followed by a FITC-conjugated mouse anti-rat IgG (RRID:AB_465229) and an AlexaFluor (AF)594-conjugated goat anti-mouse IgG (RRID: AB_2762825). Western blot for HLA-A,B,C was performed on whole cell lysates using the mAb HC10 (RRID: AB_2728622; produced in-house) and a mAb to α-tubulin (RRID: AB_1210457) as loading control, followed by a horseradish peroxidase-conjugated goat anti-mouse IgG (RRID: AB_2728714) and Pierce ECL reagents (ThermoFisher).

### Isolation and identification of HLA-I-bound peptides

Frozen cell pellets were lysed for 1 h at 4°C with 1% (w/v) octyl-b-D glucopyranoside (Sigma) diluted in 10 mM Tris-HCl pH 8.0, 150 mM NaCl, 5 mM EDTA and 0.1% (v/v) Complete Protease Inhibitor Cocktail (Roche), with vortexing every 30 min. Lysates were cleared by centrifugation and pHLA-I complexes isolated by immunoaffinity purification using W6/32 mAb (RRID:AB_1107730) crosslinked to protein A beads (GE Healthcare) with dimethyl pimelimidate (Sigma). Beads were loaded into GELoader Tips (ThermoFisher), washed and pHLA-I complexes eluted with 10% (v/v) acetic acid. The eluates were further concentrated by SpeedVac, acidified with 10% trifluoroacetic acid (TFA) and loaded into pre-washed and -equilibrated C18 stage tips (ThermoFisher). After further washing, peptides were eluted with 50% (v/v) acetonitrile (ACN) and 0.05% (v/v) TFA. Samples were dried, resuspended in 0.05% TFA 2% ACN and spiked with 10 fmol/µL of CMV pp65_495-503_ peptide (NLVPMVATV) as an internal control.

Using a Dionex UltiMate 3000 RSLCnano system, peptides were loaded onto a C18 Acclaim PepMap nanocolumn (5 mm × 300 µm; ThermoFisher) and separated with an Acclaim PepMap C18 nanocolumn (50 cm × 75 µm; Dionex) coupled to a nano-electrospray ionization Q Exactive HF mass spectrometer (ThermoFisher). Separation was achieved with a linear gradient of 2-90% buffer B (80% ACN, 0.05% formic acid) at a flow rate of 220 nL/min over 90 min at 35-45°C.

Full MS spectra were acquired from 375 to 1,500 m/z, at a resolution of 60,000 with an automatic gain control (AGC) target of 3,000,000. Precursors were selected using Top20 mode at a resolution of 15,000 with an AGC target of 50,000, a maximum injection time of 120 ms and a dynamic exclusion of 20-30 s. Unassigned and precursor ion charge states of 1 or >5 (or >4 in some cases) were excluded. The peptide match option was set to ‘preferred’.

Data was analyzed using PEAKS 8.1 (Bioinformatics Solutions). Sequence interpretation was carried out using no enzyme specification, a mass tolerance precursor of 10 ppm for peptides and 0.05 for fragment ions. Sequence matching was done against a database containing: a) all human SwissProt entries (downloaded 11/2017); b) aa sequences translated from RNA-sequencing of the CVB1 and CVB3 strains used; c) an in-house database compiled as described^15^, containing 679,576 and 530,424 predicted peptide splice products composed of major known and candidate β-cell antigen sequences with CVB1 and CVB3 sequences, respectively; d) large T antigen from simian virus (UniProt P03070) used for β-cell immortalization; and e) putative polyproteins from a xenotropic murine leukemia virus (UniProt F8UU35, F8UU37) known to be present in ECN90 β cells^41^. Post-translational modifications detected with the PEAKS PTM built-in module were disregarded as likely artifactual and the unmodified aa sequence was retained. A false discovery rate of 5% was set using a parallel decoy database search. Only the peptides with an 8-12 aa length compatible with HLA-I binding were retained.

### *In-silico* selection of candidate epitopes

A parallel *in-silico* search for nonamer peptides predicted to bind HLA-A2 or HLA-A3 was performed for all 6 CVB strains, using the following reference sequences: CVB1 AAC00531.1 (UniProt P08291), CVB2 AOW42548.1 (UniProt A0A1D8QMF3), CVB3 AFC88096.1 (UniProt H9B4F8), CVB4 AHB37371.1 (UniProt W8DN51), CVB5 AFO42818.1 (UniProt I7AVS5), CVB6 AAF12719.1 (UniProt Q9QL88). The aa positions of each peptide are reported with reference to the first serotype listed. Criteria used for this search were: a binding affinity (IC50) <40 nM, as predicted by both NetMHCpan 4.0 through the IEDB HLA-I prediction interface and NetMHC 4.0; or an identity of ≥3 consecutive aa with a β-cell antigen sequence (INS, GAD, IA-2, ZnT8, IAPP, IGRP) in central TCR contact residues (positions P3-P7). Peptides containing cysteines or found only in CVB2 or CVB6 were excluded. Two CVB peptides previously reported^42^ were appended to this list: CVB1-2-3-4-5-6_1502-1510_ (GIIYIIYKL, IEDB #20311) and CVB2-4-5-6_1589-1598_ (ILMNDQEVGV, IEDB #27205).

### HLA-I binding assays

Peptides predicted to bind HLA-A2 or -A3 (NetMHCStabPan 1.0) were synthesized (>95% pure, Synpeptide) and binding tested using biotin-tagged HLA-A2 and -A3 monomers (ImmunAware), as described^18^. Briefly, biotinylated monomers were folded (1.2 nM final concentration) and captured on streptavidin-coated beads (6-8 µm, Spherotech), followed by incubation with anti-β2-microglobulin BBM.1 mAb (RRID:AB_626748) and then with AF488-labeled goat polyclonal anti-mouse IgG (RRID:AB_2728715). Positive controls included the HLA-A2-binding peptides Flu MP_58-66_ (GILGFVFTL) and TYR_369-377_ (YMDGTMSQV) and the HLA-A3-binding peptide Flu NP_265-273_ (ILRGSVAHK). The peptide CHGA_382-390_ (HPVGEADYF) was used as negative control for both HLA-A2 and -A3. Beads were acquired on a BD LSRFortessa cytometer and analyzed by gating on single beads and AF488-positive events. Results are expressed as relative fluorescence intensities (RFIs), i.e. the median fluorescence intensity fold increase of the tested pHLA-I complex compared with the negative control. For HLA-A2, peptides with RFI>6 were considered as intermediate binders and RFI>20 as strong binders. For HLA-A3, peptides with RFI>15 were scored as intermediate binders and RFI>40 as strong binders.

### Study participants, specimens, HLA typing and CVB serology

Study participants (<u>Table S1</u>) were recruited in Paris and Miami and gave written informed consent under ethics approval DC-2015-2536 Ile-de-France I and University of Miami IRB protocol 1995-119, respectively. HLA-A2 (A*02:01) and -A3 (A*03:01) typing was performed with AmbiSolv primers (Dynal/ThermoFisher) and by DKMS (Tübingen, Germany) in case of ambiguities. Blood was drawn into 9-mL sodium heparin tubes and PBMCs isolated and frozen/thawed as described^15,43^. Frozen/thawed samples were used throughout the study. Neutralizing antibody titers against the 6 CVB serotypes were measured with a plaque neutralization assay as described^44^. Frozen PLN, spleen and blood cells, provided by nPOD and the Human Atlas of Neonatal Development and Early Life Immunity program (HANDEL-I; <u>Table S3</u>) were thawed in the presence of benzonase (50 U/ml, Merck 70664-3) and mifepristone (100 nM, Invitrogen H110-01).

### Combinatorial HLA Class I multimer assays

HLA-A2 and -A3 multimers (MMrs) (immunAware) were produced and used as described^15,18,19^. pHLA complexes were conjugated with fluorochrome-labeled streptavidins (1:4 ratio) and used at a final concentration of 8-27 nmol/L, modulated in order to correct for the variable staining index of each streptavidin and thus visualize a distinct double-MMr^+^ population for each fluorochrome pair. PBMCs were thawed in prewarmed AIM-V medium (ThermoFisher), washed, counted and incubated with 50 nmol/L dasatinib for 30 min at 37°C before negative magnetic CD8^+^ T-cell enrichment (RRID:AB_2728716). Staining with combinatorial double-coded MMr panels was performed for 20 min at 20°C in 20 µL PBS-dasatinib for 10^7^ cells followed, without washing, by staining at 4°C for 20 min with mAbs CD3-APC-H7 (RRID:AB_1645475), CD8-PE-Cy7 (RRID:AB_396852), CD45RA-FITC (RRID:AB_395879), CCR7-BV421 (RRID:AB_2728119), CD25-BV510 (RRID:AB_2629671), PD-1-BV605 (RRID:AB_2563212) and Live/Dead Aqua (ThermoFisher). After washing, cells were acquired on an LSRFortessa configured as described^15^ and analyzed with FlowJo v10 and GraphPad Prism 9. Candidate epitopes that did not yield any appreciable MMr staining or fluorochrome pairs not coding for a pHLA MMr provided negative controls for each panel. Positive control peptides included in the MMr panels were HCV PP_1406-1415_ (naïve viral control; KLVALGINAV; IEDB # 32208) and/or MelanA_26-35_ ELA (naïve self control; ELAGIGILTV; IEDB #12941) and Influenza virus (Flu) MP_58-66_ (recall viral control; GILGFVFTL; IEDB #20354) for HLA-A2; and HIV Nef_84-92_ (naïve viral control; AVDLSHFLK; IEDB #5295) and Influenza virus NP_265-273_ (recall viral control; ILRGSVAHK; IEDB #27283) for HLA-A3.

For the detection of CVB1_1132-1140_/GAD_272-280_ cross-reactive CD8^+^ T cells, the CVB1_1132-1140_ MMr pair (PE-CF594 and BV711) shared the BV711 fluorochrome with the GAD_272-280_ MMr pair (PE and BV711), thus allowing selective gating of T cells recognizing only CVB1_1132-1140_ or GAD_272-280_ (i.e., double MMr^+^) or both (i.e., triple MMr^+^), as described^19^.

### *In-situ* MMr staining on spleen, PLN and pancreas tissue sections

Frozen tissue sections of spleen, PLN and different regions of the pancreas (head, body and/or tail) were from nPOD non-diabetic (n=5), double-aAb^+^ (n=2) and T1D (n=8) cadaveric organ donors (<u>Table S4</u>; Ethics Committee #215/17S at Technical University of Munich and Helmholtz Munich Institute of Diabetes Research).

For validating *in-situ* staining with APC-labeled MMrs, PBMCs were isolated from HLA-A2^+^ and HLA-A3^+^ healthy donors using Lymphoprep density gradient centrifugation and stored at −80°C till use. Frozen/thawed PBMCs were magnetically depleted of CD8^−^ cells by MACS positive magnetic selection (Miltenyi). Isolated CD8^+^ cells were then incubated for 20 min with the indicated MMrs (10 nM) diluted in 2% goat serum in PBS and fixed with 4% Image-iT Fixative Solution (ThermoFisher) for 15 min. CD8^+^ cells were then cytospun on a microscope slide and blocked with 10% goat serum and Human TruStain FcX (RRID:AB_2818986; 1:20) for 20 min at 4°C. MMr signal amplification was obtained with a mouse anti-APC mAb (RRID:AB_345357; 1:1,000) for 1 h at room temperature (RT). An AF647-conjugated goat anti-mouse IgG1 Ab (RRID:AB_141658; 1:800) was then added together with a rabbit anti-human CD3 Ab (RRID:AB_2335677; 1:300) for 1 h at RT. CD8^+^ cells were detected by adding an AF750-conjugated goat anti-rabbit IgG Ab (RRID:AB_1500687; 1:800). Sections were counterstained with Hoechst 33342 (Invitrogen; 1:5,000) for 8 min and mounted with ProLong Gold Antifade (Invitrogen). Sections were then scanned using an Axio Scan.Z1 slide scanner (Zeiss) and a 20×/0.8NA Plan-Apochromat (a=0.55 mm) objective.

Frozen tissue sections were prefixed with 0.2% Image-iT Fixative Solution for 5 min at 4°C. After gentle washing, sections were incubated with MMrs (10 nM) diluted as above for 3 h at 4°C, washed, fixed with 4% Image-iT for 15 min, blocked with 10% goat serum and Human TruStain FcX as above, and incubated with mouse anti-APC mAb (1:50) for 1 h at RT. AF647-conjugated goat anti-mouse IgG1 (1:800) was then added for 1 h at RT together with rabbit anti-human CD8 Ab (RRID:AB_304247; 1:150), followed by AF750-conjugated goat anti-rabbit IgG (1:200). Hoechst counterstaining, mounting and scanning were performed as above.

For image analysis, a mean of 23 regions of interest (ROI) per donor from the spleen and whole slide images for PLNs were analyzed for each section. Briefly, the total number of cells per ROI or tissue area was determined using QuPath^45^ with Stardist, a deep-learning-based method of 2D nucleus detection^46^. The presence of MMr^+^CD8^+^ T cells was assessed manually. Cell density values were obtained by calculating the number of positive cells divided by the area (in mm²). Pancreatic tissue sections were first assessed for the presence of insulitis. Due to the scarcity of islet-infiltrating immune cells and MMr^+^ cells, further quantitative assessment was not performed. VP1 immunofluorescence staining was performed on pancreas tissue sections from 3 T1D donors: nPOD 6070 (RRID:SAMN15879127; female, 23-year-old, T1D duration 7 years), nPOD 6228 (RRID:SAMN15879284; male, 13-year-old; T1D duration 0 years); nPOD 6243 (RRID:SAMN15879299; male, 13-year-old, T1D duration 5 years). Four-micrometer sections were dewaxed in Histoclear, rehydrated in degrading ethanol concentrations (100%, 95%, 70%) before heat-induced epitope retrieval (10 mM citrate, pH 6). Sections were blocked with 5% goat serum and incubated with mouse anti-VP1 Ab 5-D8/1 (RRID:AB_2118128; 1:1,500) for 1 h at RT. AF488-conjugated goat anti-mouse IgG Ab (RRID:AB_2534069; 1:400) was then added for 1 h at RT. This was followed by incubation with guinea pig anti-insulin Ab (RRID:AB_2617169; 1:600) for 1 h at RT. AF555-conjugated goat anti-guinea pig IgG Ab (RRID:AB_2535856; 1:400) was then added for 1 h at RT along with DAPI (1 µg/ml). Sections were mounted and scanned using the Vectra Polaris slide scanner (Akoya Biosciences). Quantification was performed using the Indica HALO image analysis platform. Manual annotation around the cell periphery was performed for >50 VP1^+^ and VP1^−^ β cells to measure cell area, perimeter and diameter.

### *In-vitro* T-cell stimulation, TCR sequencing and re-expression

To generate PLN T-cell lines, frozen/thawed single-cell suspensions were bulk-sorted by flow cytometry for CD4^+^ and CD8^+^ T cells. CD8^+^ T cells (10^5^/well, 4 wells/each) were stimulated with anti-CD28 (5 µg/ml) and irradiated autologous spleen-derived EBV-immortalized B cells pulsed with the indicated peptides (50 µg/ml), with IL-2 (20 U/ml) and IL-15 (10 ng/ml) added on day 6. On day 14, cultures were split and recalled with 50 µg/ml of the stimulating peptide, the negative control peptide WNV_3098-3106_ or DMSO diluent pulsed on irradiated autologous B cells with anti-CD28. After 48 h, supernatants were collected and assayed for cytokine secretion by cytokine bead array (BD).

Single-cell TCR sequencing was performed on frozen/thawed cells rested overnight, followed by CD8^+^ magnetic enrichment and MMr staining, sorting in 96-well plates with a BD FACSAria II and a multiplex nested PCR, as described^17^. Sequences were assigned using IMGT (https://www.imgt.org) and visualized as heatmaps using R software (https://github.com/raivokolde/pheatmap).

5KC T-hybridoma TCR transductants were generated as described^21^. Briefly, 5KC cells were transduced with the NFAT-driven fluorescent reporter ZsGreen-1 along with human CD8 by spinoculation with retroviral supernatant produced from Phoenix-ECO cells (ATCC CRL-3214). These cells were subsequently transduced with retroviral vectors encoding a murine TRAC chimeric TCRα gene followed by a porcine Teschovirus-1 2A (P2A) peptide and a murine TRBC chimeric TCRβ gene (Twist Bioscience).

Primary CD8^+^ T-cell transductants were generated from CD8^+^ T cells magnetically enriched from fresh PBMCs and plated at 20,000 cells/well. After activation with anti-CD3 and anti-CD28 microbeads (ThermoFisher) for 48 h in AIM-V medium, cells were transduced with lentiviral vectors encoding the above chimeric TCR construct and supplemented with 40 U/ml IL-2 (Proleukin, Clinigen).

To validate antigen specificity, transductants (20,000 cells/well in 96-well U-bottom plates) were stimulated for 18 h with individual peptides added at 10-fold serial dilutions from 100 μM to 100 fM in the presence of K562 cells (50,000 cells/well) transduced with an HLA-A2 or HLA-A3 monochain construct fused with β2-microglobulin, followed by analysis of ZsGreen-1 expression (for 5KC transductants) or intracellular staining for MIP-1β (RRID:AB_357303), TNF-α (RRID:AB_398566), IL-2 (RRID:AB_2573518) and IFN-γ (RRID:AB_1272026) for primary CD8^+^ transductants.

### *In-vitro* recall and cytotoxicity assays on CVB-infected β cells

For recall assays on 5KC TCR transductants, the HLA-A2/A3^+^ ECN90 β-cell line was plated 24 h ahead at 5×10^4^/well in a 96-well flat-bottom Matrigel-coated plate in in BSA-free DMEM/F12 Advanced medium supplemented as above. It was then pulsed for 1 h with serial concentrations of CVB1_1356-1364_/CVB3_1359-1367_ peptide or left unpulsed, followed by CVB3 infection (300 MOI) for 1 h, gentle washing, and addition of transductants. After 18 h, cells were collected, stained with Live/Dead Red and fixed with the eBioscience Foxp3/Transcription Factor Staining Buffer Set. NFAT-driven ZsGreen fluorescence was then acquired on a BD LSRFortessa after gating on viable single cells.

For CVB-eGFP infection assays, ECN90 β cells were seeded at 1.5×10^4^/well (∼ 40% confluency) in pre-coated 96-well plates and incubated overnight. Following infection with CVB-eGFP^39^ at different MOIs for 1 h at 37°C, Incucyte Cytotox Red (Essen Bioscience #ESS4632; 1:4,000) was added for counting dead cells. Real-time image acquisition was performed on an Incucyte S3 (Sartorius) at 4 images/well and 1 scan/h for 48 h. Images were processed with the Incucyte software by defining masks on the green (CVB-eGFP^+^) and red (dead) β-cell areas.

For T-cell cytotoxicity assays, ECN90 β cells were stained with 1 µM CellTracker Red CMTPx dye (ThermoFisher #C34552) for 30 min in BSA-free medium and seeded as above. After 6 h, they were infected with CVB3 at different MOIs for 1 h at 37°C, or loaded with 1 µM cognate CVB peptide as positive control. Following addition of primary CD8^+^ T-cell transductants at different T:β-cell ratios, Incucyte Cytotox Green (Essen Bioscience #ESS4633; 1:4,000) was added for counting dead cells. Real-time image acquisition and processing were performed as above by defining single cell and cluster masks on the green (dead-cell) and red (β-cell) area. β-cell death was defined as the percent green/red area out of total red area in each well after subtraction of the 8 h background signal.

### IFN measurements

IFN-α2a, IFN-β, IFN-γ and IFN-λ concentrations in the supernatants (25 µl/each) of CVB-infected and non-infected ECN90 β cells, cultured alone or with primary CD8^+^ T-cell transductants, were measured by MSD U-PLEX assays (Meso Scale Discovery #K15094K-1).

### Resource availability

#### Lead contact

Further information and requests for resources and reagents should be directed to and will be fulfilled by the lead contact, Roberto Mallone.

#### Materials availability

All unique/stable reagents generated in this study are available from the Lead Contact upon reasonable request with a completed Materials Transfer Agreement.

#### Data and code availability

Immunopeptidomics datasets have been deposited under PRIDE: PXD042711. Immunofluorescence digital images and flow cytometry and real-time imaging raw files are available from the Lead Contact upon reasonable request with a completed Material Transfer Agreement. No codes were specifically generated for this work.

## SUPPLEMENTAL INFORMATION

**Table S1. Donors recruited for T-cell experiments (related to Fig. 2, 3 and S4).** Columns display study groups, donor code, demographics, HLA-A2/A3 typing, titers of neutralizing antibodies (NAbs) against CVB serotypes (positive titers of ≥16 in red, borderline titers of 4 in blue; corresponding to serum dilutions of 1/16 and 1/4, respectively; na, not available), T1D duration and the corresponding symbols used in Fig. 2, 3 and S4. HLA-A2^+^ donors recruited in Miami for T-cell validation experiments are indicated with black-filled symbols. T1D and healthy donors recruited in Miami are indicated with white- or blue-filled symbols. All other donors were recruited in Paris.

**Table S2. Summary of T-cell screening and validation experiments (related to Fig. 2 and 3).** The first columns detail the identity of each peptide. For the screening round (Fig. 2), MMr^+^CD8^+^ T-cell frequencies, MMr^+^ counts and percent effector/memory fractions are listed. For the more stringent validation round (Fig. 3), donors/peptides with <3 MMr^+^ cells counted were excluded from the final analysis, with frequencies noted as 1.0E-7 (i.e. 10^−7^) and percent effector/memory fractions not assigned (NA). Merged table rows indicate peptides with high sequence homology. Validated epitopes are displayed in red fonts.

**Table S3. Spleen, PLN and PBMC specimens analyzed by flow cytometry MMr staining (related to Fig. 4).** Results available in the nPOD repository are reported for pancreas VP1 staining, Enterovirus qPCR and CVB proteomics (prot) analyses. The last 3 sets of columns detail the number of total CD8^+^ T cells analyzed and the MMr^+^ counts and frequencies for CVB peptides, Flu peptides (Flu MP_58-66_ for HLA-A2, Flu NP_265-273_ for HLA-A3) and naïve viral peptides (HCV PP_1406-1415_ for HLA-A2, HIV nef_84-92_ for HLA-A3). HLA-A2-restricted CVB1-2-3-4-5-6_1246-1254_ and CVB1_271-279_ and HLA-A3-restricted CVB1_1356-1364_/CVB3_1359-1367_ and CVB1_1132-1140_ were analyzed. For CVB1_271-279_ and CVB1_1132-1140_, the corresponding values are marked with an asterisk. Positive results (≥3 MMr^+^ counts and ≥10^−5^ frequency) are shown in bold (and red for CVB). na, not applicable or not available. The non-diabetic donor group comprised specimens from nPOD and HANDEL repositories (HDL codes). Results obtained with the two sets of specimens were similar and were therefore pooled for figure representation.

**Table S4. nPOD spleen and PLN specimens analyzed by *in-situ* MMr staining (related to Fig. 5).** Results available in the nPOD repository are reported for pancreas VP1 staining, Enterovirus qPCR and CVB proteomics (prot) analyses. The last 2 sets of columns detail the number of MMr^+^ cells counted and densities per mm^2^ for CVB peptides, CMV/Flu peptides and naïve West Nile Virus (WNV) peptides (detailed in Fig. S11). na, not applicable or not available. For HLA-A2/A3^+^ cases 6211 and 6046, HLA-A3-restricted peptides were studied.

**Figure S1. Mapping of HLA-eluted and *in silico* predicted peptides and conservation across CVB serotypes**. ECN90 β cells were infected with either CVB1 or CVB3. The pHLA-I complexes were purified and peptides sequenced by MS (bold fonts, highlighted in yellow). A parallel *in silico* search across relevant strains (see Methods) identified nonamer peptides predicted to bind HLA-A2 or HLA-A3 (bold font, highlighted in grey). Within these peptides, the aa conserved across serotypes are highlighted in yellow (for HLA-eluted peptides) or grey (for *in silico* predicted peptides). All the peptides were tested for their ability to bind to the corresponding HLA-I monomers (Fig. S2). Binding peptides thus identified (encircled with blue dashes) were screened by combinatorial HLA-I multimer (MMr) assays. Those scored as MMr^+^ in this screening phase are encircled with red dashes. Peptides whose T-cell recognition was confirmed in validation experiments (Fig. 3, Table S2) are encircled with continuous red lines, with their aa positions indicated on top. CVB1 and CVB3 aa sequences are those translated from the RNA-sequencing of strains used for *in-vitro* infection. The aa differences between the experimental (RNA-sequenced) strains used for HLA-I peptidomics and the reference sequences used for *in-silico* predictions are indicated in red letters.

**Figure S2. Eluted vs. predicted HLA-I-binding CVB peptides, binding measurements and reproducibility of MMr assays. A.** Schematic of the CVB genome. **B.** Distribution of HLA-I-eluted vs. predicted HLA-I-binding peptides across the P1, P2 and P3 proteins. The graph shows the percent distribution of peptides expected within P1, P2 and P3 based on the aa length (first bar) and on the number of unique predicted binders of each protein (second bar; using NetMHCcons 1.1) compared to the number of unique binders experimentally eluted (third bar). Reference CVB1/CVB3 aa sequences were those translated from the RNA-sequencing of strains used for *in-vitro* infection. \*\*\**p*<0.0001 by χ^2^ test. **C.** HLA-I binding measurements. Biotinylated recombinant HLA-A2/A3 molecules were folded with peptides and β2-microglobulin. Complexes were captured on streptavidin-coated beads and stained for β2-microglobulin. Representative staining for HLA-A2 (left) and HLA-A3 complexes (right) folded with CVB peptides (blue profiles) displaying strong, intermediate and no binding. Binding controls (Flu MP_58-66_ and TYR_369-377_ for HLA-A2; Flu NP_265-273_ for HLA-A3; yellow profiles) and non-binding controls (CHGA_382-390_; red profiles) are shown for comparison. **D-E.** HLA-eluted and *in-silico* predicted peptides were tested for their binding to HLA-A2 (D) and HLA-A3 (E). Horizontal dotted lines indicate cut-off values for scoring strong, intermediate and non-binders. Bars represent mean ± SEM values from 2-3 different experiments of the relative fluorescence intensity (RFI), i.e. the median fluorescence intensity arbitrary units (A.U.) normalized to the CHGA_382-390_ non-binding peptide. *In-silico* predicted HLA-A2-binding peptides CVB1_150-158_ (KLPDALSQM; and its binding homologous ZnT8_245-253_, VMG<u>DAL</u>GSV), CVB1-2-3-4-5-6_1502-1510_ (GIIYIIYKL; IEDB #20311) and CVB1-2-3-5-6_2051-2059_ (VIASYPWPI) did not confirm as binders and are therefore not listed in Fig. 1G. **F.** Reproducibility of CVB peptide-loaded HLA-A2 MMr assays. Donor H004N was analyzed using two separate blood draws and the same MMr batch, with PBMCs stained freshly after isolation and not fixed (black circles) or stained on frozen/thawed samples fixed after staining (white circles). Coefficient of variation (CV) of measured MMr^+^CD8^+^ T-cell frequencies was 27%. **G.** Reproducibility of CVB peptide-loaded HLA-A3 MMr assays. Donor H001M was analyzed using frozen/thawed PBMCs from two separate blood draws and two separate MMr batches, with PBMCs fixed after staining. CV=21%. MMr^+^ counts, the number of total CD8^+^ T cells acquired and the percent effector/memory fraction among MMr^+^ cells are shown below each symbol. NA, not applicable (effector/memory fraction not assigned when <5 MMr^+^ cells detected).

**Figure S3. Representative staining with the HLA-A2 and HLA-A3 MMr panels used for T-cell validation experiments.** Frozen/thawed PBMCs from donor H004N (HLA-A2/A3^+^) were magnetically depleted of CD8^−^ cells before staining, acquisition, and analysis. **A.** Gating strategy for the HLA-A2 MMr panel. After verifying the stability of each fluorescence signal over the acquisition time (example shown for APC), cells were sequentially gated on singlets (FSC-A/FSC-H dot plot), live cells (Live/Dead Aqua^−^), small lymphocytes (FSC-A/SSC-A dot plot), CD3^+^CD8^+^ T cells, and total PE^+^, APC^+^, PE-CF594^+^, BV711^+^, and BV786^+^ MMr^+^ cells. Using Boolean operators, these latter gates allowed to selective visualize each double-MMr^+^ population by including only those events positive for the corresponding fluorochrome pair. The CD45RA/CCR7 staining distribution of total CD3^+^CD8^+^ T cells is also shown. **B.** The final HLA-A2 MMr readout obtained is shown for the 10 peptides analyzed. Events corresponding to each epitope-reactive population are overlaid in different colors within each dot plot. Peptides scoring positive for this donor are marked with an asterisk. Numbers in each panel indicate the MMr^+^CD8^+^ T-cell frequency out of total CD8^+^ T cells and the percent effector/memory fraction among MMr^+^ cells (NA when not assigned, i.e. <5 MMr^+^ cells counted). **C.** The final HLA-A3 MMr readout obtained is shown for the 9 peptides analyzed (with the PE/APC fluorochrome pair left empty). Data representation is the same as in panel B. **D.** The gating strategy based on positivity for 2 MMr fluorochromes precludes visualization of CVB1_1132-1140_/GAD_272-280_ cross-reactive CD8^+^ T cells, since they are positive for 3 fluorochromes (PE, PE-CF594 and the shared BV711 fluorochrome). Hence, one further gating was performed for these fractions by selecting cells positive for only 2 fluorochromes (PE-CF594/BV711^+^, PĒ, labeled as “CVB1_1132-1140_ only”; or PE/BV711^+^, PE-CF594−, labeled as “GAD_272-280_ only”) or for all 3 (PE/PE-CF594/BV711^+^, labeled as “CVB1 × GAD”). The resulting overlaid dot plot is displayed (with cells negative for all 3 fluorochromes depicted in grey), with most T cells (shown in red) cross-recognizing CVB1_1132-1140_ and GAD_272-280_ epitopes.

**Figure S4. Circulating HLA-A2-restricted CVB-reactive CD8^+^ T cells in pediatric donors. A-B.** Definition of the minimal CD8^+^ T-cell sampling size required to accurately measure CVB epitope-reactive CD8^+^ T-cell frequencies. In panel A, decreasing numbers of total CD8^+^ T cells from a single blood draw were acquired for donor H004N after staining with CVB peptide-loaded HLA-A2 MMrs. The MMr^+^CD8^+^ T-cell frequency measured on the maximal number (2×10^6^) of total CD8^+^ T cells acquired was considered as the gold standard (indicated with crossed symbols), and the CV of frequencies measured with lower numbers of acquired T cells calculated accordingly. For CVB epitopes recognized by MMr^+^CD8^+^ T cells showing a mean frequency >3/10^5^ (vertical dotted line), the acquisition of a minimal number of 100,000 total CD8^+^ T cells (red diamonds) yields an accurate frequency measurement (CV<35%; horizontal dotted line). For MMr^+^CD8^+^ T-cell frequencies <2/10^5^, the acquisition of a minimal number of 380,000 total CD8^+^ T cells (green circles) is required to maintain a CV<35%. For MMr^+^CD8^+^ T-cell frequencies <7/10^6^, the acquisition of a minimal number of 520,000 total CD8^+^ T cells (blue circles) is required to maintain a CV<35%. Panel B depicts the median frequency and distribution of MMr^+^CD8^+^ T cells using the indicated CVB peptides in pediatric donors with >500,000 total CD8^+^ T cells acquired. Based on these frequencies, the cut-offs defined in panel A for the minimal number of total CD8^+^ T cells acquired were applied for each peptide, as indicated by grey shades: ≥520,000 (blue circles) for CVB5_945-953_, ≥380,000 (green circles) for CVB1-2-4-5-6_271-279_, ≥100,000 (red diamonds) for all other epitopes. Using these cut-offs for each peptide of interest, only those samples allowing for an accurate frequency measurement were retained for final analysis. **C-D.** Frequency (C) and percent effector/memory phenotype (D) of MMr^+^CD8^+^ T cells recognizing the indicated CVB epitopes or a control memory Flu MP_58-66_ epitope in T1D (blue) and healthy (red) pediatric donors (listed in Table S3). Data representation is the same as in Fig. 3. Children recruited in Paris and Miami are represented with symbols filled in white and black, respectively. \**p*=0.04 and \*\**p*=0.01 by Mann-Whitney U test. The HLA-A2 MMr staining panel was as follows: CVB1-2-3-4-5-6_1671-1679_ PE/APC, CVB5-6_169-177_ PE/BV711, CVB5_945-953_ PE/BV786, CVB1-2-3-4-5-6_1246-1254_ APC/BV711, CVB1-2-4-5-6_271-279_ APC/BV786, Flu MP_58-66_ BV711/BV786.

**Figure S5. CVB-reactive PLN T cells in HLA-A3^+^ T1D nPOD case 6480, representative MMr staining on PLN cells and CVB *vs.* Influenza virus MMr^+^CD8^+^ T cells across tissues from nPOD donors.** A. MMr panels used for the analysis of nPOD spleen, PLN and PBMC specimens. The combined HLA-A2/HLA-A3 MMr panels were used for specimens from HLA-A2/A3^+^ nPOD donors. The homologous CVB1_1132-1140_/ GAD_272-280_ peptides are marked with an asterisk. The MMr panel for PLN and PBMC specimens was also used for splenocytes from nPOD donors 6338, 6386, 6420, 6461 and 6480. The extended phenotypic panel used for PLN and PBMC specimens is also detailed. **B-E.** PLN CD8^+^ T cells were stimulated with the indicated CVB or INS_38-46_ (INS_B14-22_) peptides for 14 days, followed by a 48-h recall with the stimulating peptide, the negative control WNV PP_3098-3106_ peptide or DMSO diluent and cytokine measurement on culture supernatants. Besides IFN-γ, TNF-α and IL-6 were also secreted in response to CVB1_1765-1774_ peptide (not shown). Data is depicted as mean ± SEM of triplicate wells. \**p*≤0.02, \*\**p*=0.003, \*\*\**p*=0.0007 by paired Student’s t test. **F.** Representative HLA-A3 MMr staining on PLN cells from T1D nPOD case 6362. Given the limited cell numbers available, frozen-thawed PLN cells were directly stained and acquired, without prior magnetic depletion of CD8^−^ cells. The gating strategy is the same as in Fig. S3A, but each fluorochrome pair was here used for only one MMr, in order to allow sorting of individual CVB double-MMr^+^ cells for TCR sequencing, with control Flu MMrs labeled with a single BV711 fluorochrome. The final readout obtained is shown for the 3 peptides analyzed. Each dot plot displays a color-coded overlay of individual double-MMr^+^ subsets to visualize the separation of each epitope-reactive CD8^+^ T-cell fraction relative to the MMr− population (gray). Numbers in each panel indicate the MMr^+^CD8^+^ T-cell frequency out of total CD8^+^ T cells and the percent effector/memory fraction among MMr^+^ cells. PD-1^+^ cells (not shown) were gated on MMr^+^ effector/memory fractions. **G-H.** Influenza virus MMr^+^CD8^+^ T cells in lymphoid tissues from nPOD cases (see details in Table S3). MMr^+^CD8^+^ T-cell frequencies (G) and percent PD-1^+^CD25^−^ MMr^+^ cells (H; for donors/epitopes with cell counts ≥5) are depicted across available tissues. **I.** FlowSOM uniform manifold approximation and projection (UMAP) analysis of CVB (CVB1_271-279_ and CVB1_1356-1364_) and Influenza virus (Flu MP_58-66_ and Flu NP_265-273_) MMr^+^CD8^+^ cells from all available tissues. Left, clustering of the two populations; right, the projected surface PD-1 expression.

**Figure S6. Expanded TCR CDR3β clonotypes in individual CVB1/CVB3_1356-1364/1359-1367_-reactive CD8^+^ T cells from the blood of living CVB-seropositive healthy donors and common CDR3 motifs. A.** Expanded clonotypes were defined as those found in at least 2 MMr^+^ cells in the same donor, and are clustered according to frequency (one columns per donor) and sequence similarity (one row per CDR3β clonotype). **B.** Sequence logo plots displaying CDR3β (top) and CDR3α (bottom) motifs among all TCRs sequenced from CVB1/CVB3_1356-1364_/_1359-1367_ MMr^+^CD8^+^ T cells isolated from tissues and blood. The x-axis shows the CDR3 aa position after MMseqs2 centroid alignment. The y-axis shows the information content, with the size of each aa symbol proportional to its frequency. **C.** Distribution of *TRBV* (top), *TRBJ* (middle) and *TRAV* (bottom) gene usage among 378 TCRs sequenced from CVB1/CVB3_1356-1364_/_1359-1367_ MMr^+^CD8^+^ T cells isolated from tissues and blood.

**Figure S7. Validation of *in-situ* MMr staining. A.** MMrs used for *in-situ* staining on human tissues. **B.** Representative immunofluorescence images of isolated CD8^+^ T cells cytospun on microscope glass slides. Cells were stained with the indicated pooled CMV/Flu, pooled CVB or single WNV peptide-loaded MMrs (red) and CD3 (green). Cell nuclei are stained in blue. White arrows point to MMr^+^CD3^+^ cells. Scale bars: 20 μm. **C.** Correlation between CVB MMr^+^CD8^+^ cell frequencies measured by flow cytometry and CVB MMr^+^CD8^+^ cell densities quantified by tissue immunofluorescence. The samples analyzed by both techniques available for this comparison were all from spleen (round symbols), barring a PLN sample from double-aAb^+^ case 6197 (square symbol).

**Figure S8. Correlation between CVB-mediated and T-cell-mediated β-cell killing. A.** r, 95% confidence interval (CI) and *p* values of the correlation between CVB-mediated and T-cell-mediated β-cell killing depicted in Fig. 7D. **B.** Equivalent inhibition of CVB-mediated single β-cell death by both transduced (T) and non-transduced (NT) CD8^+^ T cells at a 2:1 T:β-cell ratio. **C.** T-cell-mediated cluster β-cell death in the same conditions. **D.** CVB peptide-specific β-cell killing by transduced (T) and non-transduced (NT) CD8^+^ T cells. No killing is observed with non-transduced T cells.

**Figure S9. Alignment of the main CD8^+^ T-cell epitopes identified across CVB and Poliovirus serotypes.** Amino-acid differences are shaded in grey. For Poliovirus, the 3 serotypes included in the inactivated poliovirus vaccine were used as reference sequences: serotype 1 (Mahoney; GenBank V01149.1), 2 (MEF1; GenBank AY238473.1) and 3 (Saukett; GenBank KP247597.1). The alignment refers to the serotype with the closest match.

**Movie S1. Kinetics of CVB infection and CVB-eGFP transfer through filopodia in ECN90 β cells.** Real-time imaging of ECN90 β cells infected with CVB-eGFP at MOI 100 and stained with Cytotox Red to visualize dead cells. The movie shows an intact β cell in contact with the filopodia of an infected eGFP^+^ cell, turning eGFP^+^ at and subsequently protruding filopodia before dying (Cytotox Red^+^). Infected cells are labeled in green, dead cells are labeled in red.

**Movie S2. Death kinetics and morphology in CVB-infected ECN90 β cells and definition of a single-cell analysis mask.** β cells (infected at 300 MOI, in the absence of T cells) are labeled in red, dead cells are labeled in green, merged images of dead β cells are labeled in yellow. Contours in magenta red indicate the areas of single-cell death defined by the imaging software based on the preset analysis mask.

**Movie S3. Death kinetics and morphology of ECN90 β cells co-cultured with CVB-reactive CD8^+^ T-cell transductants and definition of a cell cluster analysis mask.** β cells (pulsed with 1 µM CVB1_1356-1364_ peptide, without infection) are labeled in red, dead cells are labeled in green, merged images of dead β cells are labeled in yellow. Contours in magenta red indicate the areas of clustered cell death defined by the imaging software based on the preset analysis mask.

**Movie S4A-B. Mask analyses for counting of single-cell and cluster cell death areas of CVB-infected ECN90 β cells co-cultured with CVB-reactive CD8^+^ T-cell transductants.** β cells (infected at 300 MOI and co-cultured at a 1:1 T:β-cell ratio) are labeled in red, dead cells are labeled in green, merged images of dead β cells are labeled in yellow. The analysis of the same field with the defined single-cell (A) and cluster cell analysis mask (B) is shown, with contours in magenta red indicating the areas counted with each mask.

