## Supplemental Information for "Coxsackievirus infection induces direct pancreatic β-cell killing but poor anti-viral CD8+ T-cell responses"

**Table S1. Donors recruited for T-cell experiments (related to Fig. 2, 3 and S4).** Columns display study groups, donor code, demographics, HLA-A2/A3 typing, titers of neutralizing antibodies (NAbs) against CVB serotypes (positive titers of ≥16 in red, borderline titers of 4 in blue; corresponding to serum dilutions of 1/16 and 1/4, respectively; na, not available), T1D duration and the corresponding symbols used in Fig. 2, 3 and S4. HLA-A2$^+$ donors recruited in Miami for T-cell validation experiments are indicated with black-filled symbols. T1D and healthy donors recruited in Miami are indicated with white- or blue-filled symbols. All other donors were recruited in Paris.

| Study group | Code | Age (yrs) | Gender (M/F) | HLA | CVB1 NAbs | CVB2 NAbs | CVB3 NAbs | CVB4 NAbs | CVB5 NAbs | CVB6 NAbs | T1D duration (yrs) | Symbols |
|---|---|---|---|---|---|---|---|---|---|---|---|---|
| Screen (n=9) | H079O | 38 | F | A2 | ≥16 | ≥16 | 0 | 0 | ≥16 | 4 | na | ○ |
| | H110S | 60 | F | A3 | 0 | ≥16 | 0 | ≥16 | ≥16 | 4 | na | ◇ |
| | H112S | 42 | F | A3 | ≥16 | ≥16 | 0 | ≥16 | 4 | ≥6 | na | □ |
| | H158O | 56 | M | A2 | ≥16 | ≥16 | ≥16 | ≥16 | ≥16 | ≥16 | na | □ |
| | H170S | 40 | F | A2 | ≥16 | ≥16 | ≥16 | ≥16 | ≥16 | 0 | na | ▽ |
| | H458C | 33 | M | A3 | ≥16 | ≥16 | ≥16 | ≥16 | ≥16 | 4 | na | △ |
| | H460C | 24 | M | A2 | 0 | 0 | ≥16 | ≥16 | ≥16 | 0 | na | ◉ |
| | H463C | 29 | F | A2 | ≥16 | 4 | ≥16 | 0 | ≥16 | 4 | na | ▣ |
| | H477C | 34 | F | A3 | 4 | ≥16 | ≥16 | ≥16 | ≥16 | 4 | na | ▽ |
| Screen + Validation (n=6) | H004N | 45 | M | A2/A3 | 0 | ≥16 | ≥16 | 0 | ≥16 | 4 | na | ◈ |
| | H106S | 48 | F | A2 | ≥16 | ≥16 | ≥16 | ≥16 | ≥16 | ≥16 | na | ◇ |
| | H455C | 25 | F | A2 | ≥16 | ≥16 | ≥16 | 0 | 4 | ≥16 | na | △ |
| | H466C | 25 | F | A2 | ≥16 | ≥16 | ≥16 | ≥16 | ≥16 | ≥16 | na | ▽ |
| | H478C | 36 | F | A3 | 0 | ≥16 | 0 | ≥16 | ≥16 | 0 | na | ◉ |
| | H487C | 39 | M | A2 | na | na | na | na | na | na | na | △ |
| Validation (n=10) | H001M | 27 | M | A3 | na | na | na | na | na | na | na | ● |
| | H002M | 36 | M | A3 | na | na | na | na | na | na | na | ◆ |
| | H003M | 29 | M | A3 | na | na | na | na | na | na | na | ■ |
| | H004M | 23 | M | A3 | na | na | na | na | na | na | na | ▲ |
| | H005M | 28 | M | A3 | na | na | na | na | na | na | na | ▼ |
| | H006M | 33 | M | A3 | na | na | na | na | na | na | na | ⊗ |
| | H372C | 26 | F | A2 | 0 | ≥16 | ≥16 | ≥16 | 0 | 4 | na | ⊗ |
| | H423C | 26 | M | A3 | na | na | na | na | na | na | na | ○ |
| | H459C | 26 | F | A3 | ≥16 | ≥16 | ≥16 | 0 | 4 | 4 | na | ▣ |
| | H494C | 56 | F | A2/A3 | na | na | na | na | na | na | na | ⊗ |
| T1D (n=11) | B001M | 8 | M | A2 | ≥16 | ≥16 | 0 | ≥16 | 0 | 4 | 3 | ◇ |
| | B002M | 12 | M | A2 | 0 | 0 | ≥16 | ≥16 | 0 | 0 | 4 | □ |
| | B003M | 12 | M | A2 | 4 | ≥16 | 0 | 0 | 0 | 0 | 7 | ○ |
| | B004M | 19 | M | A2 | 0 | 4 | 0 | ≥16 | ≥16 | 4 | 0.6 | △ |
| | B005M | 16 | M | A2 | 4 | ≥16 | ≥16 | 0 | ≥16 | 0 | 0.6 | ▽ |
| | B006M | 9 | F | A2 | 4 | 4 | 0 | ≥16 | ≥16 | 4 | 6 | ⊗ |
| | B091S | 7 | M | A2 | na | na | na | na | na | na | 0 | ● |
| | B241R | 14 | M | A2 | ≥16 | 0 | ≥16 | 0 | 0 | 4 | 0.02 | ◆ |
| | B244V | 17 | M | A2 | 0 | 0 | 0 | ≥16 | ≥16 | 4 | 0 | ■ |
| | B252R | 11 | M | A2 | ≥16 | ≥16 | ≥16 | ≥16 | ≥16 | ≥16 | 0 | ▲ |
| | B234R | 11 | M | A2 | 0 | ≥16 | ≥16 | ≥16 | ≥16 | 4 | 0.02 | ▼ |
| | | **12 (7-19)** | **91% M / 9% F** | | **60%** | **70%** | **50%** | **70%** | **60%** | **70%** | **0.6 (0-7)** | |
| Healthy (n=10) | HB001M | 9 | M | A2 | na | na | na | na | na | na | na | ◇ |
| | HB002M | 18 | M | A2 | ≥16 | 0 | ≥16 | ≥16 | 0 | 4 | na | □ |
| | HB003M | 13 | F | A2 | ≥16 | ≥16 | ≥16 | ≥16 | 4 | 4 | na | ○ |
| | HB004M | 14 | M | A2 | 4 | ≥16 | 0 | 4 | 4 | 4 | na | △ |
| | HB005M | 11 | M | A2 | 0 | ≥16 | ≥16 | ≥16 | 0 | 0 | na | ▽ |
| | HB006M | 9 | F | A2 | 4 | 0 | ≥16 | 0 | ≥16 | 4 | na | ⊗ |
| | HB007M | 16 | M | A2 | 4 | 0 | ≥16 | 0 | ≥16 | 4 | na | ◉ |
| | HB030J | 8 | M | A2 | ≥16 | ≥16 | ≥16 | ≥16 | ≥16 | ≥16 | na | ● |
| | HB052J | 13 | M | A2 | 4 | ≥16 | 0 | ≥16 | 0 | 0 | na | ◆ |
| | HB055J | 13 | M | A2 | ≥16 | 0 | 0 | ≥16 | ≥16 | ≥16 | na | ■ |
| | | **13 (8-18)** | **80% M / 20% F** | | **89%** | **50%** | **60%** | **70%** | **60%** | **70%** | | |

**Table S2. Summary of T-cell screening and validation experiments (related to Fig. 2 and 3).** The first columns detail the identity of each peptide. For the screening round (Fig. 2), MMr$^{+}$CD8$^{+}$ T-cell frequencies, MMr$^{+}$ counts and percent effector/memory fractions are listed. For the more stringent validation round (Fig. 3), donors/peptides with <3 MMr$^{+}$ cells counted were excluded from the final analysis, with frequencies noted as 1.0E-7 (i.e. $10^{-7}$) and percent effector/memory fractions not assigned (NA). Merged table rows indicate peptides with high sequence homology. Validated epitopes are displayed in red fonts.

| HLA | Strain | Position | Sequence | Pipeline | Screening round | | | Validation round | |
|---|---|---|---|---|---|---|---|---|---|
| | | | | | Frequency median | MMr+ counts median (range) | % effector/memory median (range) | Frequency median | % effector/memory median (range) |
| HLA-A2 n=24 n=5 | CVB3 | 235-244 | KLVQRVVYNA | MS | 1.6 E-6 | 1 (0-3) | 0 (0-0) | | |
| | CVB3 | 1038-1048 | SILEKSLKALV | MS | 1.2 E-6 | 1 (0-2) | 0 (0-100) | | |
| | CVB1/CVB3 | 1359-1366 | KINMPMSV | MS | 1.0 E-7 | 0 (0-1) | 0 (0-0) | | |
| | CVB1-2-5 | 60-68 | IMIKSMPAL | In silico | 2.2 E-5 | 30 (8-155) | 26 (0-70) | 1.0 E-7 | NA |
| | CVB4-6 | 60-68 | VMIKSLPAL | In silico | 5.3 E-6 | 2 (0-5) | 20 (20-20) | | |
| | CVB5 | 169-177 | YLGRAGYTV | In silico | 5.4 E-5 | 14 (6-50) | 10 (4-55) | 7.3 E-6 | 14 (0-33) |
| | CVB1-2-4-5-6 | 271-279 | IVMPYINSV | In silico | 2.2 E-6 | 2 (0-24) | 85 (0-100) | 2.1 E-5 | 43 (26-82) |
| | CVB2 | 291-299 | FTLMIIPFV | In silico | 1.5 E-6 | 1 (0-3) | 0 (0-100) | | |
| | CVB1-2-3-4-5-6 | 456-464 | AMATGKFLL | In silico | 1.0 E-7 | 0 (0-1) | 0 (0-0) | | |
| | CVB1-5 | 555-563 | RMLKDTPFI | In silico | 1.0 E-7 | 0 (0-1) | 0 (0-0) | | |
| | CVB2-5 | 711-719 | VLTHQIMYV | In silico | 1.0 E-7 | 0 (0-0) | 0 (0-0) | | |
| | CVB4 | 943-951 | YQSHVLLAV | In silico | 1.9 E-6 | 2 (0-5) | 0 (0-100) | | |
| | CVB5 | 945-953 | YQTHVLLAV | In silico | 5.6 E-5 | 25 (9-47) | 24 (7-55) | 6.7 E-6 | 8 (0-55) |
| | CVB1-2-3-4-5-6 | 1246-1254 | KLNSSVYSL | In silico | 3.0 E-5 | 13 (1-36) | 33 (0-70) | 1.3 E-5 | 38 (0-83) |
| | CVB1-2-3-4-5-6 | 1289-1297 | QMVSSVDFV | In silico | 2.7 E-6 | 1 (0-5) | 40 (40-40) | | |
| | CVB1-2-3-4-6 | 1307-1315 | GILFTSPFV | In silico | 1.0 E-7 | 0 (0-3) | 0 (0-33) | | |
| | CVB5 | 1311-1319 | FLFTSPFVL | In silico | 8.0 E-6 | 4 (1-14) | 29 (20-56) | | |
| | CVB1-2-3-4-5-6 | 1509-1517 | KLFAGFQGA | In silico | 1.0 E-7 | 0 (0-0) | 0 (0-0) | | |
| | CVB1-3 | 1585-1593 | LMNDQEIGV | In silico | 7.4 E-7 | 1 (0-12) | 50 (0-100) | | |
| | CVB2-4-5-6 | 1589-1598 | ILMNDQEVGV | In silico | 2.5 E-5 | 11 (1-892) | 40 (24-99) | 1.0 E-7 | NA |
| | CVB1-5-6 | 1626-1634 | FLAKEEVEV | In silico | 1.0 E-7 | 0 (0-1) | 0 (0-0) | | |
| | CVB2-4 | 1631-1639 | FLAREEAEV | In silico | 5.3 E-6 | 2 (0-10) | 34 (25-43) | | |
| | CVB1-2-3-4-5-6 | 1671-1679 | RMLMYNFPT | In silico | 2.8 E-5 | 24 (0-154) | 25 (0-67) | 6.2 E-6 | 16 (0-33) |
| | CVB1-2-3-4-5-6 | 2107-2115 | FLVHPVMPM | In silico | 1.0 E-7 | 0 (0-0) | 0 (0-0) | | |
| HLA-A3 n=12 n=4 | CVB1 | 804-813 | RIYFKPKHVK | MS | 4.3 E-5 | 16 (10-67) | 66 (23-88) | 1.0 E-7 | NA |
| | CVB1 | 1132-1140 | KILPEVKEK | MS | 2.0 E-5 | 9 (4-21) | 58 (20-71) | | |
| | *CVB1 only* | *1132-1140* | *KILPEVKEK* | *MS* | | | | *1.0 E-7* | *24 (20-29)* |
| | *GAD only* | *272-280* | *KMFPEVKEK* | *MS* | | | | *1.0 E-7* | *23 (20-55)* |
| | CVB1 x GAD | | | | | | | 1.3 E-5 | 60 (23-100) |
| | CVB1/CVB3 | 1233-1241/1236-1244 | SVATNLIGR | MS | 5.4 E-6 | 2 (1-6) | 42 (0-83) | | |
| | CVB1/CVB3 | 1356-1364/1359-1367 | KINMPMSVK | MS | 1.0 E-5 | 4 (0-89) | 95 (80-100) | 3.1 E-5 | 98 (80-100) |
| | CVB2 | 1361-1369 | KVNMPMSVK | In silico | 1.9 E-5 | 7 (0-15) | 61 (40-100) | | |
| | CVB3 | 1519-1528 | GAYTGVPNQK | MS | 1.0 E-7 | 0 (0-2) | 0 (0-0) | | |
| | CVB3 | 1765-1774 | VLRSGDPRLK | MS | 1.4 E-5 | 6 (0-15) | 68 (50-75) | 1.8 E-5 | 78 (56-92) |
| | CVB5 | 760-768 | SMFYDGWAK | In silico | 1.9 E-5 | 12 (2-25) | 33 (0-68) | 2.9 E-5 | 33 (9-94) |
| | CVB1-2-3-4-5-6 | 1204-1212 | KMSNYIQFK | In silico | 1.0 E-7 | 0 (0-0) | 0 (0-0) | | |
| | CVB1-2-3-4-5-6 | 2024-2032 | IIIRTLMLK | In silico | 4.4 E-6 | 2 (1-3) | 0 (0-0) | | |
| | CVB1-2-3-4-5-6 | 2027-2035 | RTLMLKVYK | In silico | 3.0 E-6 | 1 (0-6) | 25 (16-33) | | |
| | CVB2-3-4-5 | 2161-2169 | KIRSVPVGR | In silico | 1.0 E-7 | 0 (0-0) | 0 (0-0) | | |

**Table S3. Spleen, PLN and PBMC specimens analyzed by flow cytometry MMr staining (related to Fig. 4).** Results available in the nPOD repository are reported for pancreas VP1 staining, Enterovirus qPCR and CVB proteomics (prot) analyses. The last 3 sets of columns detail the number of total $CD8^+$ T cells analyzed and the $MMr^+$ counts and frequencies for CVB peptides, Flu peptides (Flu $MP_{58-66}$ for HLA-A2, Flu $NP_{265-273}$ for HLA-A3) and naïve viral peptides (HCV $PP_{1406-1415}$ for HLA-A2, HIV $nef_{84-92}$ for HLA-A3). HLA-A2-restricted CVB1-2-3-4-5-$6_{1246-1254}$ and $CVB1_{271-279}$ and HLA-A3-restricted $CVB1_{1356-1364}$/$CVB3_{1359-1367}$ and $CVB1_{1132-1140}$ were analyzed. For $CVB1_{271-279}$ and $CVB1_{1132-1140}$, the corresponding values are marked with an asterisk. Positive results (≥3 $MMr^+$ counts and $\geq 10^{-5}$ frequency) are shown in bold (and red for CVB). na, not applicable or not available. The non-diabetic donor group comprised specimens from nPOD and HANDEL repositories (HDL codes). Results obtained with the two sets of specimens were similar and were therefore pooled for figure representation.

| | nPOD case RRID SAMN# | Sex | Age (yrs) | T1D (yrs) | Positive aAbs | Pancreas VP1/qPCR/ Prot | HLA | Spleen | | | | | | | PLNs | | | | | PBMCs | | | | |
|---|---|---|---|---|---|---|---|---|---|---|---|---|---|---|---|---|---|---|---|---|---|---|---|---|
| | | | | | | | | CD8+ | CVB MMr+ | | Flu MMr+ | | HCV/HIV MMr+ | | CD8+ | CVB MMr+ | | Flu MMr+ | | CD8+ | CVB MMr+ | | Flu MMr+ | |
| | | | | | | | | # ×10³ | # | Freq | # | Freq | # | Freq | # ×10-3 | # | Freq | # | Freq | # ×10-3 | # | Freq | # | Freq |
| T1D (n=13) | 6046-15879103 | F | 19 | 8 | IA-2/ZnT8 | +/+/+ | A2<br>A3 | 173.0 | 2<br>8 | 1.2e-5<br>4.6e-5 | na<br>33 | na<br>1.9e-4 | na<br>1 | na<br>5.8e-6 | na | na | na | na | na | na | na | na | na | na |
| | 6052-15879109 | M | 12 | 1 | IA/IA-2 | +/+/− | A2 | 86.3 | 1 | 1.2e-5 | 0 | 0 | 0 | 0 | na | na | na | na | na | na | na | na | na | na |
| | 6195-15879251 | M | 19 | 5 | IA/GAD/IA-2/ZnT8 | −/+/+ | A3 | 73.3 | 1 | 1.4e-5 | 12 | 1.6e-4 | 0 | 0 | na | na | na | na | na | na | na | na | na | na |
| | 6211-15879267 | F | 24 | 4 | IA/GAD/IA-2/ZnT8 | +/+/+ | A2<br>A3 | na | na | na | na | na | na | na | 38.1 | 20*<br>1 | 5.4e-4*<br>2.7e-5 | 8 | 2.2e-4 | 46.7 | 14*<br>1 | 3.1e-4*<br>2.2e-5 | 14 | 3.1e-4 |
| | 6212-15879268 | M | 20 | 5 | IA | +/+/− | A2 | 47.6 | 0 | 0 | 8 | 1.7e-4 | 1 | 2.1e-5 | na | na | na | na | na | na | na | na | na | na |
| | 6224-15879280 | F | 21 | 2 | None | −/na/na | A2 | 30.2 | 0 | 0 | 3 | 1.0e-4 | 0 | 0 | na | na | na | na | na | na | na | na | na | na |
| | 6243-15879299 | M | 13 | 5 | IA | +/na/na | A2 | 122.8 | 0 | 0 | 43 | 3.5e-4 | 2 | 1.6e-5 | na | na | na | na | na | na | na | na | na | na |
| | 6265-15879319 | M | 11 | 8 | IA/GAD | −/−/na | A3 | 126.2 | 13 | 1.0e-4 | 1 | 7.9e-6 | 1 | 7.9e-6 | na | na | na | na | na | na | na | na | na | na |
| | 6325-15879379 | F | 20 | 6 | IA/GAD/IA-2 | +/−/+ | A2 | 128.4 | 1 | 7.8e-6 | 2 | 1.6e-5 | 0 | 0 | na | na | na | na | na | na | na | na | na | na |
| | 6362-15879415 | M | 25 | 0 | GAD | +/+/+ | A3 | 40.5 | 6 | 1.5e-4 | 17 | 4.2e-4 | 0 | 0 | 125.7 | 29<br>25* | 2.3e-4<br>2.0e-4* | 54 | 4.3e-4 | na | na | na | na | na |
| | 6438-15879491 | F | 39 | 10 | GAD | na | A2 | 220.2 | 3 | 1.4e-5 | 166 | 7.5e-4 | 0 | 0 | na | na | na | na | na | na | na | na | na | na |
| | 6456-15879509 | F | 30 | 0 | GAD/ZnT8 | −/na/na | A2 | 118.2 | 0 | 0 | 22 | 1.9e-4 | 0 | 0 | na | na | na | na | na | na | na | na | na | na |
| | 6480-15879533 | M | 17 | 3 | IA/IA-2 | na | A3 | 17.5 | 3 | 1.7e-4 | 0 | 0 | 3 | 1.7e-4 | na | na | na | na | na | na | na | na | na | na |
| aAb+ (n=10) | 6080-15879137 | F | 69 | na | IAA/GAD | +/+/− | A2 | 9.6 | 0 | 0 | 1 | 1.0e-4 | 2 | 2.1e-4 | na | na | na | na | na | na | na | na | na | na |
| | 6151-15879207 | M | 30 | na | GAD | −/−/− | A2 | 97.4 | 1 | 1.0e-5 | 10 | 1.0e-4 | 0 | 0 | na | na | na | na | na | na | na | na | na | na |
| | 6156-15879212 | M | 40 | na | GAD | −/+/na | A3 | 45.3 | 0 | 0 | 13 | 2.9e-4 | 1 | 2.2e-5 | na | na | na | na | na | na | na | na | na | na |
| | 6158-15879214 | M | 40 | na | IAA/GAD | +/+/− | A3 | 15.1 | 0 | 0 | 0 | 0 | 0 | 0 | na | na | na | na | na | na | na | na | na | na |
| | 6171-15879227 | F | 4 | na | GAD | −/−/na | A2 | 35.1 | 4 | 1.1e-4 | 0 | 0 | 0 | 0 | 87.3 | 1<br>11* | 1.2e-5<br>1.3e-4* | 32 | 3.7e-4 | 19.3 | 9<br>5* | 5.0e-4<br>2.8e-4* | 42 | 2.2e-3 |
| | 6181-15879237 | M | 32 | na | GAD | −/+/na | A3 | 70.0 | 0 | 0 | 61 | 8.7e-4 | 1 | 1.4e-5 | na | na | na | na | na | na | na | na | na | na |
| | 6197-15879253 | M | 22 | na | GAD/IA-2 | +/+/− | A2 | 87.1 | 8 | 9.2e-5 | 2 | 2.3e-5 | 1 | 1.1e-5 | 38.1 | 0<br>25* | 0<br>6.8e-4 | 7 | 1.9e-4 | na | na | na | na | na |
| | 6347-15879401 | M | 9 | na | IAA | na | A2 | 97.4 | 1 | 1.0e-5 | 5 | 5.1e-5 | 1 | 1.0e-5 | na | na | na | na | na | na | na | na | na | na |
| | 6421-15879474 | M | 7 | na | GAD | +/−/na | A2<br>A3 | 129.8 | 2<br>2 | 1.5e-5<br>1.5e-5 | na<br>2 | na<br>1.5e-5 | na<br>2 | na<br>1.5e-5 | na | na | na | na | na | na | na | na | na | na |
| | 6429-15879482 | M | 22 | na | IAA/GAD | −/−/+ | A2 | 41.4 | 0 | 0 | 1 | 2.4e-5 | 0 | 0 | na | na | na | na | na | na | na | na | na | na |
| ND (n=15) | 6102-15879159 | F | 45 | na | None | +/−/− | A3 | 19.6 | 0 | 0 | 20 | 1.0e-3 | 0 | 0 | na | na | na | na | na | na | na | na | na | na |
| | 6140-15879197 | M | 38 | na | None | +/−/− | A3 | 17.0 | 0 | 0 | 1 | 5.9e-5 | 4 | 2.4e-4 | na | na | na | na | na | na | na | na | na | na |
| | 6338-15879392 | M | 17 | na | None | na | A2<br>A3 | 134.8 | 9*<br>0 | 6.7e-5*<br>0 | 5 | 3.8e-5 | na | na | 173.3 | 3*<br>2 | 1.7e-5*<br>1.2e-5 | 3 | 1.7e-5 | na | na | na | na | na |
| | 6368-15879421 | M | 38 | na | None | +/na/na | A3 | 307.7 | 10 | 3.2e-5 | 972 | 3.2e-3 | 2 | 6.5e-6 | na | na | na | na | na | na | na | na | na | na |
| | 6386-15879439 | M | 14 | na | None | na/na/− | A2<br>A3 | 330.2 | 48*<br>4 | 1.5e-4<br>1.2e-5 | 59 | 1.8e-4 | na | na | 674.0 | 54*<br>9 | 8.1e-5*<br>1.4e-5 | 74 | 1.1e-4 | na | na | na | na | na |
| | 6420-15879473 | M | 11 | na | None | na | A2<br>A3 | 394.3 | 18*<br>1 | 4.6e-5*<br>2.6e-6 | 19 | 4.9e-5 | na | na | na | na | na | na | na | 120.0 | 9*<br>0 | 7.6e-5*<br>0 | 7 | 5.9e-5 |

| | | | | | | | | | | | | | | | | | | | | | | | | |
|---|---|---|---|---|---|---|---|---|---|---|---|---|---|---|---|---|---|---|---|---|---|---|---|---|
| | 6461-15879514 | M | 14 | na | None | na | A2<br>A3 | 75.0 | 44*<br>3 | 6.0e-4*<br>4.1e-5 | 9 | 1.2e-4 | na | na | 37.3 | 33*<br>2 | 9.1e-4*<br>5.5e-5 | 11 | 3.0e-4 | na | na | na | na | na |
| | 6488-15879541 | F | 5 | na | None | na | A2<br>A3 | 540.0 | 1<br>23 | 1.9e-6<br>4.4e-5 | 4 | 7.6e-6 | na | na | na | na | na | na | na | na | na | na | na | na |
| | HDL_2 | F | 7 | na | None | na | A2 | 189.1 | 6<br>3 | 3.3e-5*<br>1.7e-5 | 3 | 1.7e-5 | na | na | na | na | na | na | na | na | na | na | na | na |
| | HDL_4 | M | 2 | na | None | na | A2 | 77.2 | 1<br>0 | 1.3e-5*<br>0 | 1 | 1.3e-5 | na | na | na | na | na | na | na | na | na | na | na | na |
| | HDL_5 | M | 5 | na | None | na | A2 | 108.1 | 5<br>11 | 4.8e-5*<br>1.1e-4 | 2 | 1.9e-5 | na | na | na | na | na | na | na | na | na | na | na | na |
| | HDL_19 | M | 7 | na | None | na | A2 | 106.7 | 7<br>1 | 6.9e-5*<br>9.9e-6 | 2 | 2.0e-5 | na | na | na | na | na | na | na | na | na | na | na | na |
| | HDL_24 | M | 2 | na | None | na | A2 | 40.8 | 7<br>0 | 1.8e-4*<br>0 | 1 | 2.6e-5 | na | na | na | na | na | na | na | na | na | na | na | na |
| | HDL_25 | F | 2 | na | None | na | A2<br>A3 | 59.8 | 1<br>4 | 17e-5*<br>6.9e-5 | 11 | 1.9e-4 | na | na | na | na | na | na | na | na | na | na | na | na |
| | HDL_27 | F | 0.2 | na | None | na | A2 | 64.1 | 10<br>0 | 1.7e-4*<br>0 | 0 | 0 | na | na | na | na | na | na | na | na | na | na | na | na |

**Table S4. nPOD spleen and PLN specimens analyzed by *in-situ* MMr staining (related to Fig. 5).** Results available in the nPOD repository are reported for pancreas VP1 staining, Enterovirus qPCR and CVB proteomics (prot) analyses. The last 2 sets of columns detail the number of $MMr^+$ cells counted and densities per $mm^2$ for CVB peptides, CMV/Flu peptides and naïve West Nile Virus (WNV) peptides (detailed in Fig. S7). na, not applicable or not available. For HLA-$A2/A3^+$ cases 6211 and 6046, HLA-A3-restricted peptides were studied.

| | nPOD case RRID SAMN# | Sex | Age (yrs) | T1D (yrs) | Positive aAbs | Pancreas VP1/qPCR/ Prot | HLA | Spleen | | | | | | PLNs | | | | | |
|---|---|---|---|---|---|---|---|---|---|---|---|---|---|---|---|---|---|---|---|
| | | | | | | | | CVB MMr+ | | CMV/Flu MMr+ | | WNV MMr+ | | CVB MMr+ | | CMV/Flu MMr+ | | WNV MMr+ | |
| | | | | | | | | # | Density | # | Density | # | Density | # | Density | # | Density | # | Density |
| T1D (n=8) | 6046-15879103 | F | 19 | 8 | IA-2/ZnT8 | +/+/+ | A2<br>A3 | 64 | 1.40 | 35 | 0.70 | 18 | 0.94 | 45 | 8.97 | 29 | 6.08 | 19 | 4.21 |
| | 6052-15879109 | M | 12 | 1 | IA/IA-2 | +/+/− | A2 | 134 | 3.18 | 93 | 1.46 | 11 | 0.19 | 182 | 14.46 | 130 | 10.88 | 25 | 1.99 |
| | 6211-15879267 | F | 24 | 4 | IA/GAD/IA-2/ZnT8 | +/+/+ | A2<br>A3 | 85 | 1.37 | 78 | 1.36 | 25 | 0.88 | 133 | 3.22 | 215 | 6.17 | 58 | 1.43 |
| | 6224-15879280 | F | 21 | 2 | None | −/na/na | A2 | 165 | 1.77 | 43 | 0.40 | 4 | 0.03 | 35 | 4.16 | 43 | 5.07 | 5 | 0.56 |
| | 6265-15879319 | M | 11 | 8 | IA/GAD | −/−/na | A3 | 128 | 0.83 | 78 | 0.64 | 13 | 0.42 | 17 | 2.92 | 48 | 8.75 | 11 | 1.95 |
| | 6325-15879379 | F | 20 | 6 | IA/GAD/IA-2 | +/-/+ | A2 | 83 | 1.92 | 35 | 0.55 | 19 | 0.32 | 188 | 7.80 | 57 | 4.64 | 18 | 1.40 |
| | 6438-15879491 | F | 39 | 10 | GAD | na | A2 | 153 | 3.70 | 99 | 1.72 | 4 | 0.07 | 84 | 12.80 | 82 | 5.57 | 25 | 0.54 |
| | 6480-15879533 | M | 17 | 3 | IA/IA-2 | na | A3 | 9 | 0.44 | 13 | 0.30 | 7 | 0.37 | 49 | 2.95 | 46 | 2.91 | 10 | 0.76 |
| AAb+ (n=2) | 6158-15879214 | M | 40 | na | IA/GAD | +/+/− | A3 | 159 | 13.81 | 40 | 0.79 | 14 | 0.23 | 67 | 8.13 | 70 | 4.18 | 20 | 1.12 |
| | 6197-15879253 | M | 22 | na | GAD/IA-2 | +/+/− | A2 | 48 | 5.89 | 54 | 4.30 | 11 | 0.48 | 157 | 20.30 | 145 | 22.91 | 56 | 7.92 |
| ND (n=5) | 6102-15879159 | F | 45 | na | None | +/−/− | A3 | na | na | 18 | 0.38 | 10 | 0.34 | 55 | 3.73 | 32 | 2.25 | 18 | 1.08 |
| | 6227-15879283 | F | 17 | na | None | +/na/+ | A2<br>A3 | 53<br>36 | 1.20<br>0.50 | 45<br>74 | 0.89<br>0.64 | 4<br>12 | 0.04<br>0.35 | 99<br>16 | 19.18<br>3.02 | 55<br>132 | 10.13<br>23.06 | 13<br>37 | 7.24<br>2.55 |
| | 6271-15879325 | M | 17 | na | None | −/na/− | A2 | 82 | 4.36 | 53 | 2.04 | 10 | 0.20 | 3 | 5.97 | 45 | 8.55 | 5 | 1.05 |
| | 6232-15879288 | F | 14 | na | None | +/na/na | A2 | 66 | 1.75 | 16 | 0.26 | 6 | 0.09 | 66 | 4.61 | 24 | 1.93 | 6 | 0.55 |
| | 6368-15879421 | M | 38 | na | None | +/na/na | A3 | 20 | 0.53 | 22 | 0.26 | 9 | 0.45 | 166 | 10.79 | 110 | 7.93 | 71 | 5.20 |

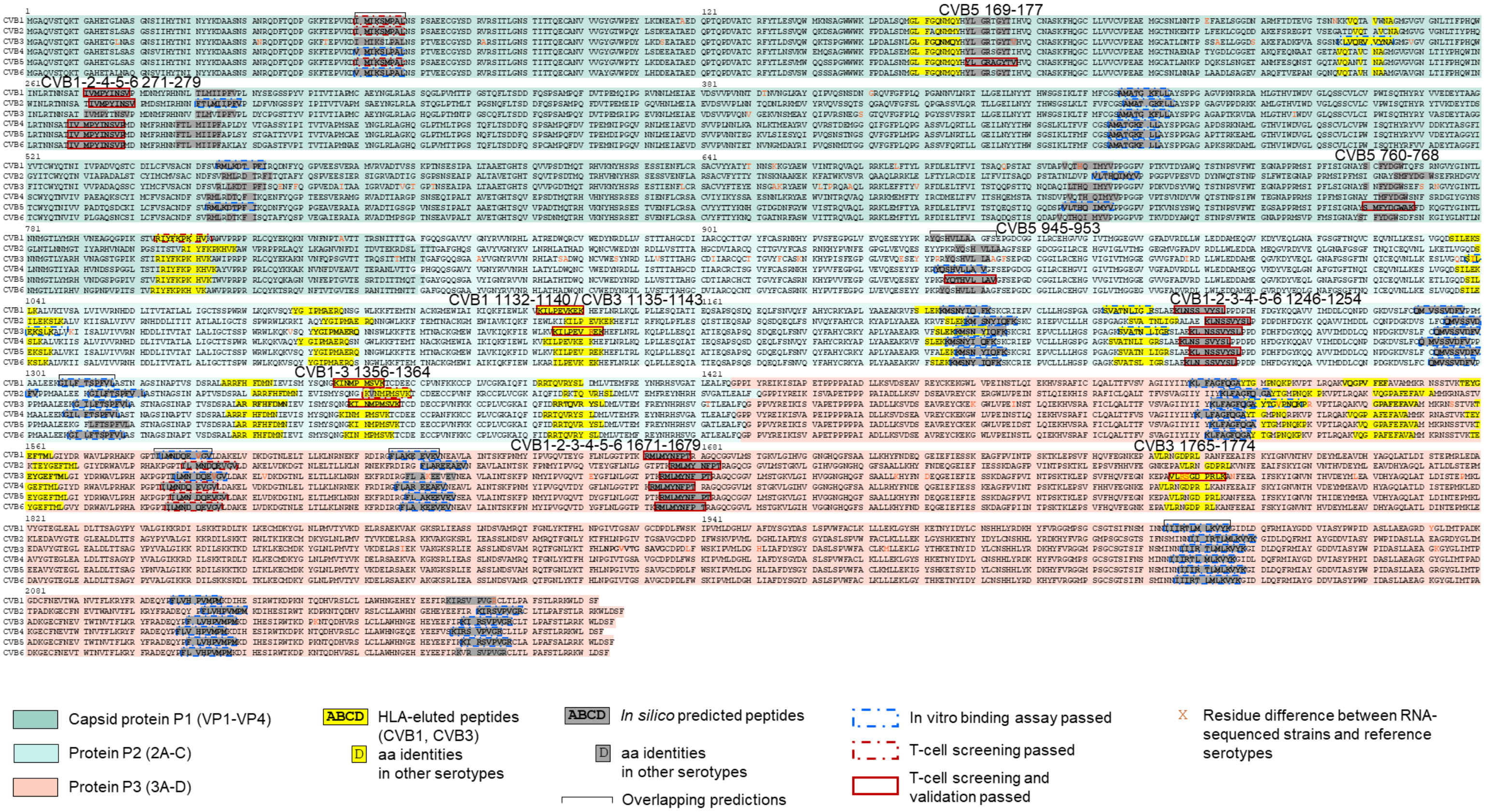

**Figure S1. Mapping of HLA-eluted and *in silico* predicted peptides and conservation across CVB serotypes**. ECN90 β cells were infected with either CVB1 or CVB3. The pHLA-I complexes were purified and peptides sequenced by MS (bold fonts, highlighted in yellow). A parallel *in silico* search across relevant strains (see Methods) identified nonamer peptides predicted to bind HLA-A2 or HLA-A3 (bold font, highlighted in grey). Within these peptides, the aa conserved across serotypes are highlighted in yellow (for HLA-eluted peptides) or grey (for *in silico* predicted peptides). All the peptides were tested for their ability to bind to the corresponding HLA-I monomers (Fig. S2). Binding peptides thus identified (encircled with blue dashes) were screened by combinatorial HLA-I multimer (MMr) assays. Those scored as MMr$^{+}$ in this screening phase are encircled with red dashes. Peptides whose T-cell recognition was confirmed in validation experiments (Fig. 3, Table S2) are encircled with continuous red lines, with their aa positions indicated on top. CVB1 and CVB3 aa sequences are those translated from the RNA-sequencing of strains used for *in-vitro* infection. The aa differences between the experimental (RNA-sequenced) strains used for HLA-I peptidomics and the reference sequences used for *in-silico* predictions are indicated in red letters.

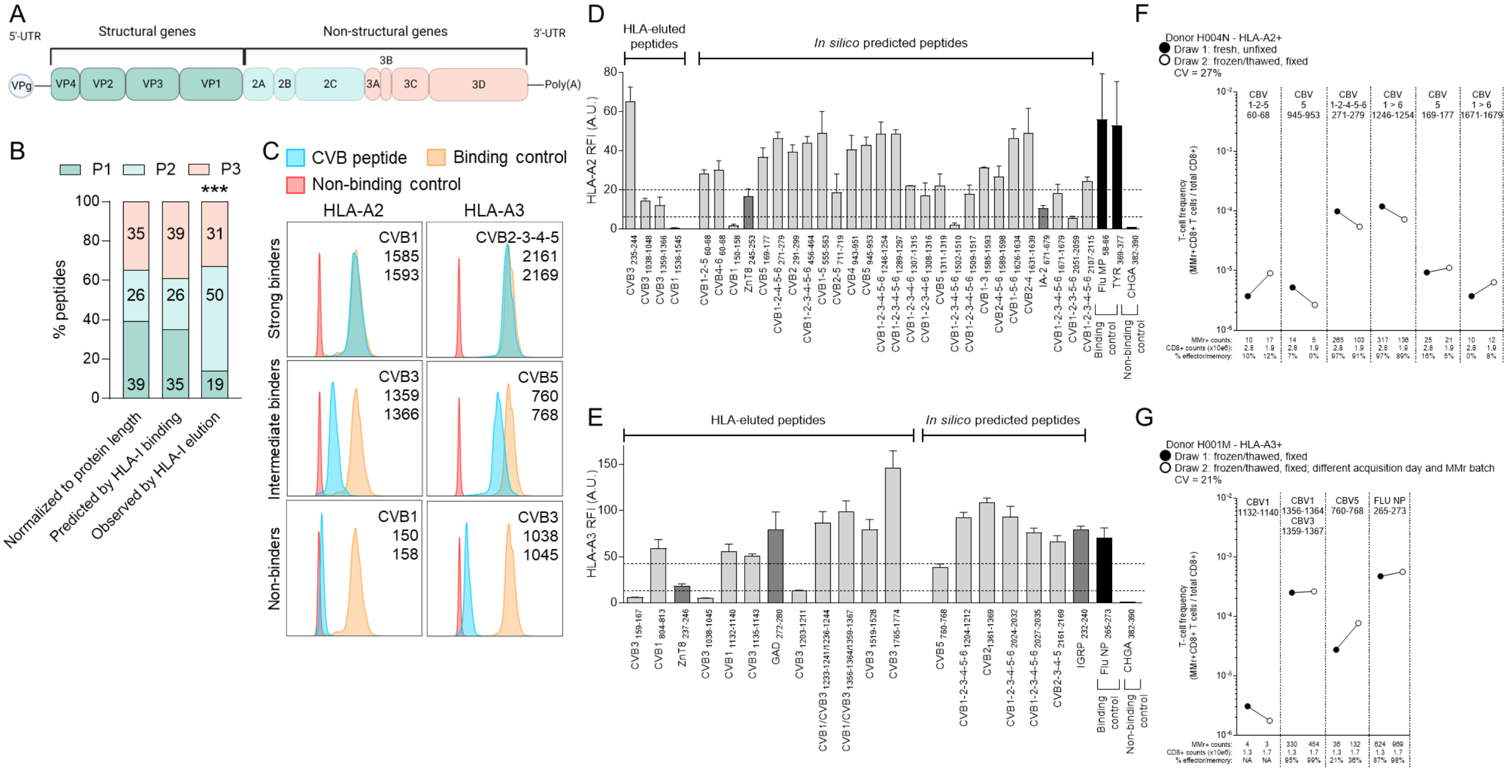

**Figure S2. Eluted vs. predicted HLA-I-binding CVB peptides, binding measurements and reproducibility of MMr assays. A.** Schematic of the CVB genome. **B.** Distribution of HLA-I-eluted vs. predicted HLA-I-binding peptides across the P1, P2 and P3 proteins. The graph shows the percent distribution of peptides expected within P1, P2 and P3 based on the aa length (first bar) and on the number of unique predicted binders of each protein (second bar; using NetMHCcons 1.1) compared to the number of unique binders experimentally eluted (third bar). Reference CVB1/CVB3 aa sequences were those translated from the RNA-sequencing of strains used for *in-vitro* infection. ***$p$<0.0001 by $\chi^2$ test. **C.** HLA-I binding measurements. Biotinylated recombinant HLA-A2/A3 molecules were folded with peptides and β2-microglobulin. Complexes were captured on streptavidin-coated beads and stained for β2-microglobulin. Representative staining for HLA-A2 (left) and HLA-A3 complexes (right) folded with CVB peptides (blue profiles) displaying strong, intermediate and no binding. Binding controls (Flu MP$_{58\text{-}66}$ and TYR$_{369\text{-}377}$ for HLA-A2; Flu NP$_{265\text{-}273}$ for HLA-A3; yellow profiles) and non-binding controls (CHGA$_{382\text{-}390}$; red profiles) are shown for comparison. **D-E.** HLA-eluted and *in-silico* predicted peptides were tested for their binding to HLA-A2 (D) and HLA-A3 (E). Horizontal dotted lines indicate cut-off values for scoring strong, intermediate and non-binders. Bars represent mean ± SEM values from 2-3 different experiments of the relative fluorescence intensity (RFI), i.e. the median fluorescence intensity arbitrary units (A.U.) normalized to the CHGA$_{382\text{-}390}$ non-binding peptide. *In-silico* predicted HLA-A2-binding peptides CVB1$_{150\text{-}158}$ (KLPDALSQM; and its binding homologous ZnT8$_{245\text{-}253}$, VMGDALGSV), CVB1-2-3-4-5-6$_{1502\text{-}1510}$ (GIIYIIYKL; IEDB #20311) and CVB1-2-3-5-6$_{2051\text{-}2059}$ (VIASYPWPI) did not confirm as binders and are therefore not listed in Fig. 1G. **F.** Reproducibility of CVB peptide-loaded HLA-A2 MMr assays. Donor H004N was analyzed using two separate blood draws and the same MMr batch, with PBMCs stained freshly after isolation and not fixed (black circles) or stained on frozen/thawed samples fixed after staining (white circles). Coefficient of variation (CV) of measured MMr$^+$CD8$^+$ T-cell frequencies was 27%. **G.** Reproducibility of CVB peptide-loaded HLA-A3 MMr assays. Donor H001M was analyzed using frozen/thawed PBMCs from two separate blood draws and two separate MMr batches, with PBMCs fixed after staining. CV=21%. MMr$^+$ counts, the number of total CD8$^+$ T cells acquired and the percent effector/memory fraction among MMr$^+$ cells are shown below each symbol. NA, not applicable (effector/memory fraction not assigned when <5 MMr$^+$ cells detected).

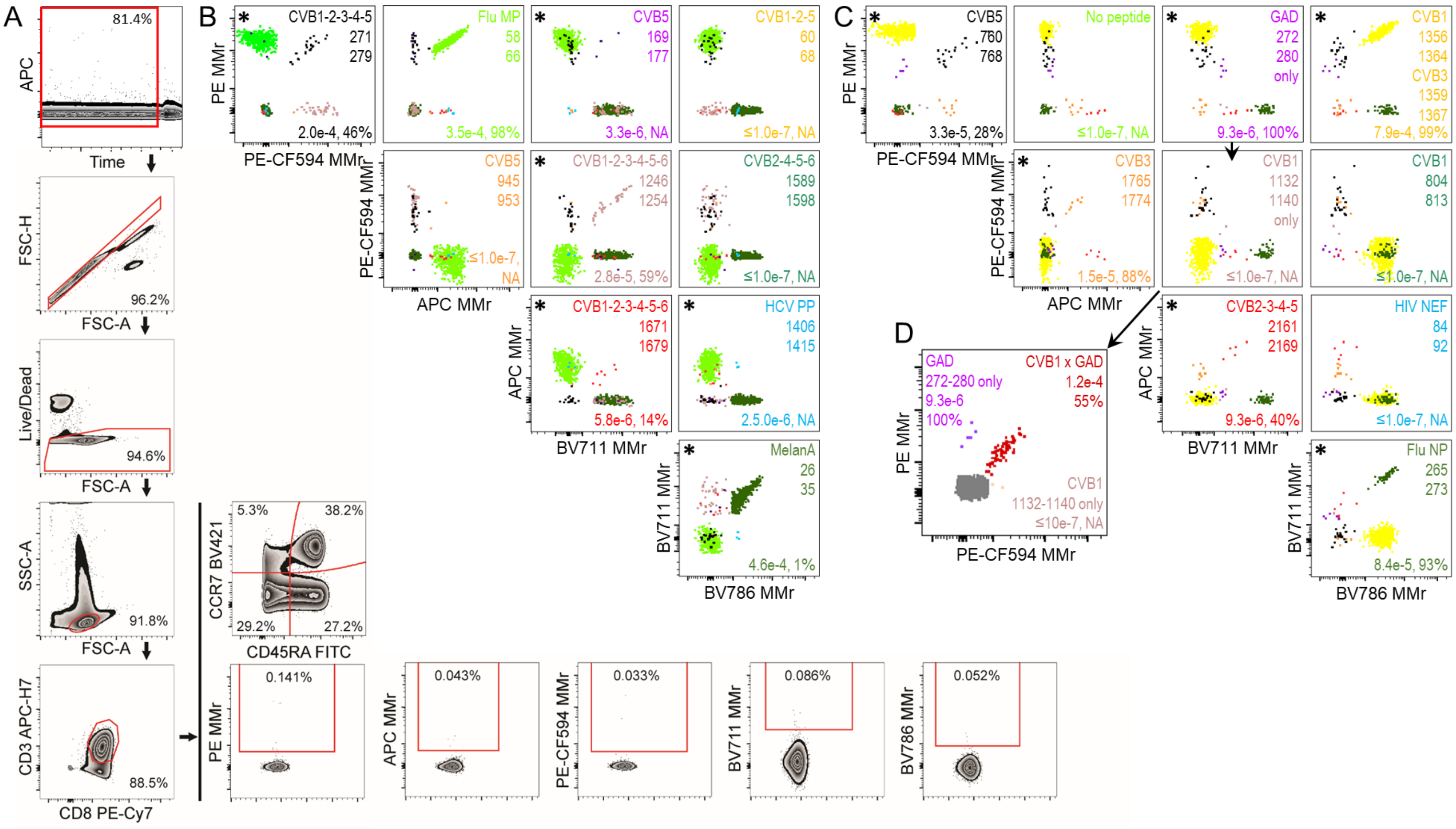

**Figure S3. Representative staining with the HLA-A2 and HLA-A3 MMr panels used for T-cell validation experiments.** Frozen/thawed PBMCs from donor H004N (HLA-A2/A3$^{+}$) were magnetically depleted of $CD8^{-}$ cells before staining, acquisition, and analysis. **A.** Gating strategy for the HLA-A2 MMr panel. After verifying the stability of each fluorescence signal over the acquisition time (example shown for APC), cells were sequentially gated on singlets (FSC-A/FSC-H dot plot), live cells (Live/Dead Aqua$^{-}$), small lymphocytes (FSC-A/SSC-A dot plot), $CD3^{+}CD8^{+}$ T cells, and total $PE^{+}$, $APC^{+}$, PE-$CF594^{+}$, $BV711^{+}$, and $BV786^{+}$ $MMr^{+}$ cells. Using Boolean operators, these latter gates allowed to selective visualize each double-$MMr^{+}$ population by including only those events positive for the corresponding fluorochrome pair. The CD45RA/CCR7 staining distribution of total $CD3^{+}CD8^{+}$ T cells is also shown. **B.** The final HLA-A2 MMr readout obtained is shown for the 10 peptides analyzed. Events corresponding to each epitope-reactive population are overlaid in different colors within each dot plot. Peptides scoring positive for this donor are marked with an asterisk. Numbers in each panel indicate the $MMr^{+}CD8^{+}$ T-cell frequency out of total $CD8^{+}$ T cells and the percent effector/memory fraction among $MMr^{+}$ cells (NA when not assigned, i.e. <5 $MMr^{+}$ cells counted). **C.** The final HLA-A3 MMr readout obtained is shown for the 9 peptides analyzed (with the PE/APC fluorochrome pair left empty). Data representation is the same as in panel B. **D.** The gating strategy based on positivity for 2 MMr fluorochromes precludes visualization of $CVB1_{1132\text{-}1140}$/$GAD_{272\text{-}280}$ cross-reactive $CD8^{+}$ T cells, since they are positive for 3 fluorochromes (PE, PE-CF594 and the shared BV711 fluorochrome). Hence, one further gating was performed for these fractions by selecting cells positive for only 2 fluorochromes (PE-CF594/$BV711^{+}$, $PE^{-}$, labeled as "$CVB1_{1132\text{-}1140}$ only"; or PE/$BV711^{+}$, PE-$CF594^{-}$, labeled as "$GAD_{272\text{-}280}$ only") or for all 3 (PE/PE-CF594/$BV711^{+}$, labeled as "CVB1 × GAD"). The resulting overlaid dot plot is displayed (with cells negative for all 3 fluorochromes depicted in grey), with most T cells (shown in red) cross-recognizing $CVB1_{1132\text{-}1140}$ and $GAD_{272\text{-}280}$ epitopes.

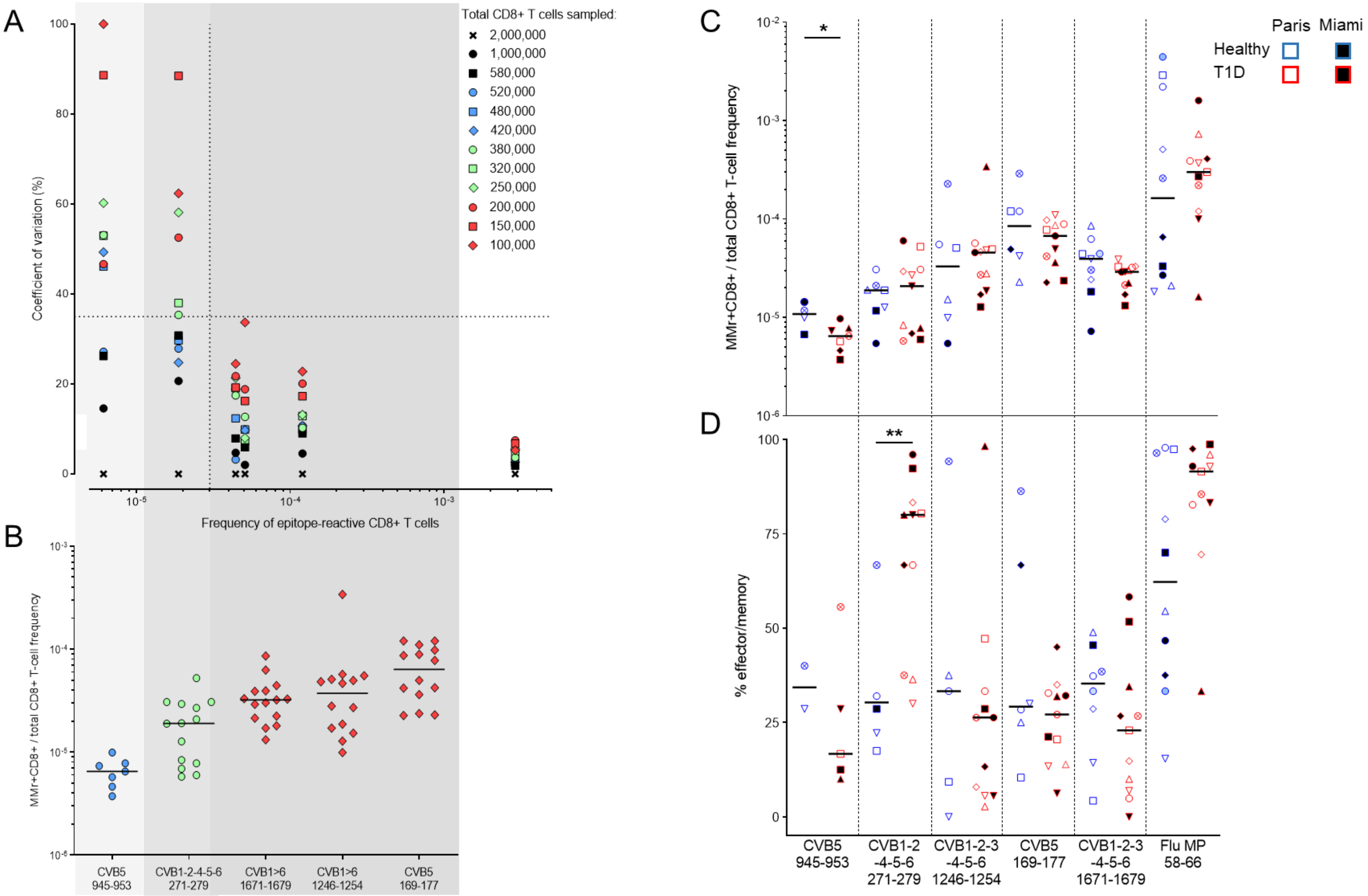

**Figure S4. Circulating HLA-A2-restricted CVB-reactive CD8$^+$ T cells in pediatric donors. A-B.** Definition of the minimal CD8$^+$ T-cell sampling size required to accurately measure CVB epitope-reactive CD8$^+$ T-cell frequencies. In panel A, decreasing numbers of total CD8$^+$ T cells from a single blood draw were acquired for donor H004N after staining with CVB peptide-loaded HLA-A2 MMrs. The MMr$^+$CD8$^+$ T-cell frequency measured on the maximal number ($2\times10^6$) of total CD8$^+$ T cells acquired was considered as the gold standard (indicated with crossed symbols), and the CV of frequencies measured with lower numbers of acquired T cells calculated accordingly. For CVB epitopes recognized by MMr$^+$CD8$^+$ T cells showing a mean frequency $>3/10^5$ (vertical dotted line), the acquisition of a minimal number of 100,000 total CD8$^+$ T cells (red diamonds) yields an accurate frequency measurement (CV<35%; horizontal dotted line). For MMr$^+$CD8$^+$ T-cell frequencies $<2/10^5$, the acquisition of a minimal number of 380,000 total CD8$^+$ T cells (green circles) is required to maintain a CV<35%. For MMr$^+$CD8$^+$ T-cell frequencies $<7/10^6$, the acquisition of a minimal number of 520,000 total CD8$^+$ T cells (blue circles) is required to maintain a CV<35%. Panel B depicts the median frequency and distribution of MMr$^+$CD8$^+$ T cells using the indicated CVB peptides in pediatric donors with >500,000 total CD8$^+$ T cells acquired. Based on these frequencies, the cut-offs defined in panel A for the minimal number of total CD8$^+$ T cells acquired were applied for each peptide, as indicated by grey shades: ≥520,000 (blue circles) for CVB5$_{945\text{-}953}$, ≥380,000 (green circles) for CVB1-2-4-5-6$_{271\text{-}279}$, ≥100,000 (red diamonds) for all other epitopes. Using these cut-offs for each peptide of interest, only those samples allowing for an accurate frequency measurement were retained for final analysis. **C-D.** Frequency (C) and percent effector/memory phenotype (D) of MMr$^+$CD8$^+$ T cells recognizing the indicated CVB epitopes or a control memory Flu MP$_{58\text{-}66}$ epitope in T1D (blue) and healthy (red) pediatric donors (listed in Table S1). Data representation is the same as in Fig. 3. Children recruited in Paris and Miami are represented with symbols filled in white and black, respectively. **p*=0.04 and ***p*=0.01 by Mann-Whitney U test. The HLA-A2 MMr staining panel was as follows: CVB1-2-3-4-5-6$_{1671\text{-}1679}$ PE/APC, CVB5-6$_{169\text{-}177}$ PE/BV711, CVB5$_{945\text{-}953}$ PE/BV786, CVB1-2-3-4-5-6$_{1246\text{-}1254}$ APC/BV711, CVB1-2-4-5-6$_{271\text{-}279}$ APC/BV786, Flu MP$_{58\text{-}66}$ BV711/BV786.

A

| MMr panel | Fluorochrome pair | Peptide |
|---|---|---|
| **Spleen specimens** (Fig. 4A-B) | | |
| **MMr panel** | **Fluorochrome pair** | **Peptide** |
| HLA-A2 | PE / APC | HCV PP 1406-1415 |
| | PE / BV711 | CVB5 169-177 |
| | PE / BV786 | CVB5 945-953 |
| | APC / BV711 | CVB1-2-3-4-5-6 1246-1254 |
| | APC / BV786 | CVB1-2-4-5-6 271-279 |
| | BV711 / BV786 | Flu MP 58-66 |
| HLA-A3 | PE / APC | HIV nef 84-92 |
| | PE / BV711 | GAD 272-280* |
| | PE / BV786 | CVB1 1356-1364 / CVB3 1359-1367 |
| | APC / BV711 | CVB1 1132-1140* |
| | APC / BV786 | CVB5 760-768 |
| | BV711 / BV786 | Flu NP 265-273 |
| HLA-A2 + HLA-A3 | PE / APC | HLA-A3 HIV nef 84-92 |
| | PE / BV711 | HLA-A3 GAD 272-280 |
| | PE / BV786 | HLA-A3 CVB1 1356-1364 / CVB3 1359-1367 |
| | APC / BV711 | HLA-A2 CVB1-2-3-4-5-6 1246-1254 |
| | APC / BV786 | HLA-A2 CVB1-2-4-5-6 271-279 |
| | BV711 / BV786 | HLA-A3 Flu NP 265-273 |
| **PLN and PBMC specimens** (Fig. 4C-D-E) | | |
| **MMr panel** | **Fluorochrome pair** | **Peptide** |
| HLA-A2 | PE/BV650 | CVB1-2-3-4-5-6 1246-1254 |
| | APC/PE-CF594 | CVB1 271-279 |
| | BV711 | Flu MP 58-66 |
| HLA-A3 | PE/BV650 | CVB1 1356-1364 / CVB3 1359-1367 |
| | APC/PE-CF594 | CVB1 1132-1140 |
| | BV711 | Flu NP 265-273 |
| HLA-A2 + HLA-A3 | PE/BV650 | HLA-A2 CVB1 271-279 |
| | APC/PE-CF594 | HLA-A3 CVB1 1356-1364 / CVB3 1359-1367 |
| | BV711 | HLA-A3 Flu NP 265-273 |
| Phenotype markers | Far Red | Live/Dead |
| | APC-H7 | CD3 |
| | PE-Cy7 | CD8 |
| | FITC | CD45RA |
| | BV421 | CCR7 |
| | BV510 | CD25 |
| | BV605 | PD-1 |

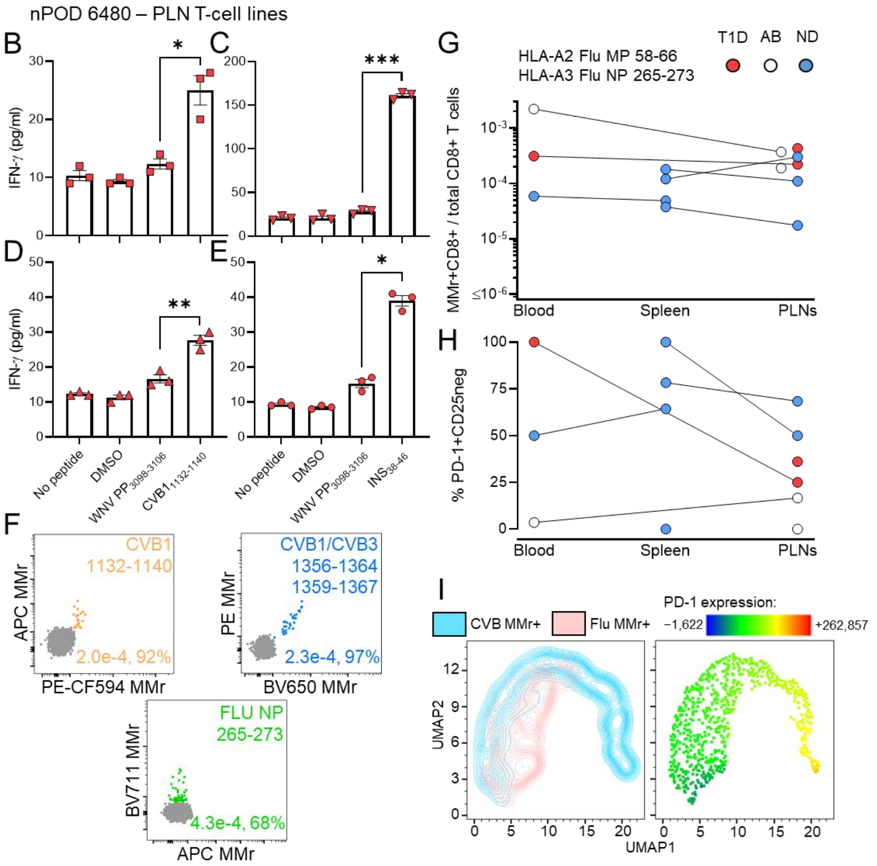

**Figure S5. CVB-reactive PLN T cells in HLA-A3$^{+}$ T1D nPOD case 6480, representative MMr staining on PLN cells and CVB *vs.* Influenza virus MMr$^{+}$CD8$^{+}$ T cells across tissues from nPOD donors. A.** MMr panels used for the analysis of nPOD spleen, PLN and PBMC specimens. The combined HLA-A2/HLA-A3 MMr panels were used for specimens from HLA-A2/A3$^{+}$ nPOD donors. The homologous CVB1$_{1132\text{-}1140}$/ GAD$_{272\text{-}280}$ peptides are marked with an asterisk. The MMr panel for PLN and PBMC specimens was also used for splenocytes from nPOD donors 6338, 6386, 6420, 6461 and 6480. The extended phenotypic panel used for PLN and PBMC specimens is also detailed. **B-E.** PLN CD8$^{+}$ T cells were stimulated with the indicated CVB or INS$_{38\text{-}46}$ (INS$_{B14\text{-}22}$) peptides for 14 days, followed by a 48-h recall with the stimulating peptide, the negative control WNV PP$_{3098\text{-}3106}$ peptide or DMSO diluent and cytokine measurement on culture supernatants. Besides IFN-γ, TNF-α and IL-6 were also secreted in response to CVB1$_{1765\text{-}1774}$ peptide (not shown). Data is depicted as mean ± SEM of triplicate wells. *$p$≤0.02, **$p$=0.003, ***$p$=0.0007 by paired Student's t test. **F.** Representative HLA-A3 MMr staining on PLN cells from T1D nPOD case 6362. Given the limited cell numbers available, frozen-thawed PLN cells were directly stained and acquired, without prior magnetic depletion of CD8$^{-}$ cells. The gating strategy is the same as in Fig. S3A, but each fluorochrome pair was here used for only one MMr, in order to allow sorting of individual CVB double-MMr$^{+}$ cells for TCR sequencing, with control Flu MMrs labeled with a single BV711 fluorochrome. The final readout obtained is shown for the 3 peptides analyzed. Each dot plot displays a color-coded overlay of individual double-MMr$^{+}$ subsets to visualize the separation of each epitope-reactive $CD8^+$ T-cell fraction relative to the $MMr^-$ population (gray). Numbers in each panel indicate the $MMr^+CD8^+$ T-cell frequency out of total $CD8^+$ T cells and the percent effector/memory fraction among $MMr^+$ cells. **G-H.** Influenza virus $MMr^+CD8^+$ T cells in lymphoid tissues from nPOD cases (see details in Table S3). $MMr^+CD8^+$ T-cell frequencies (G) and percent $PD\text{-}1^+CD25^-$ $MMr^+$ cells (H; for donors/epitopes with cell counts ≥5) are depicted across available tissues. **I.** FlowSOM uniform manifold approximation and projection (UMAP) analysis of CVB ($CVB1_{271\text{-}279}$ and $CVB1_{1356\text{-}1364}$) and Influenza virus (Flu $MP_{58\text{-}66}$ and Flu $NP_{265\text{-}273}$) $MMr^+CD8^+$ cells from all available tissues. Left, clustering of the two populations; right, the projected surface PD-1 expression.

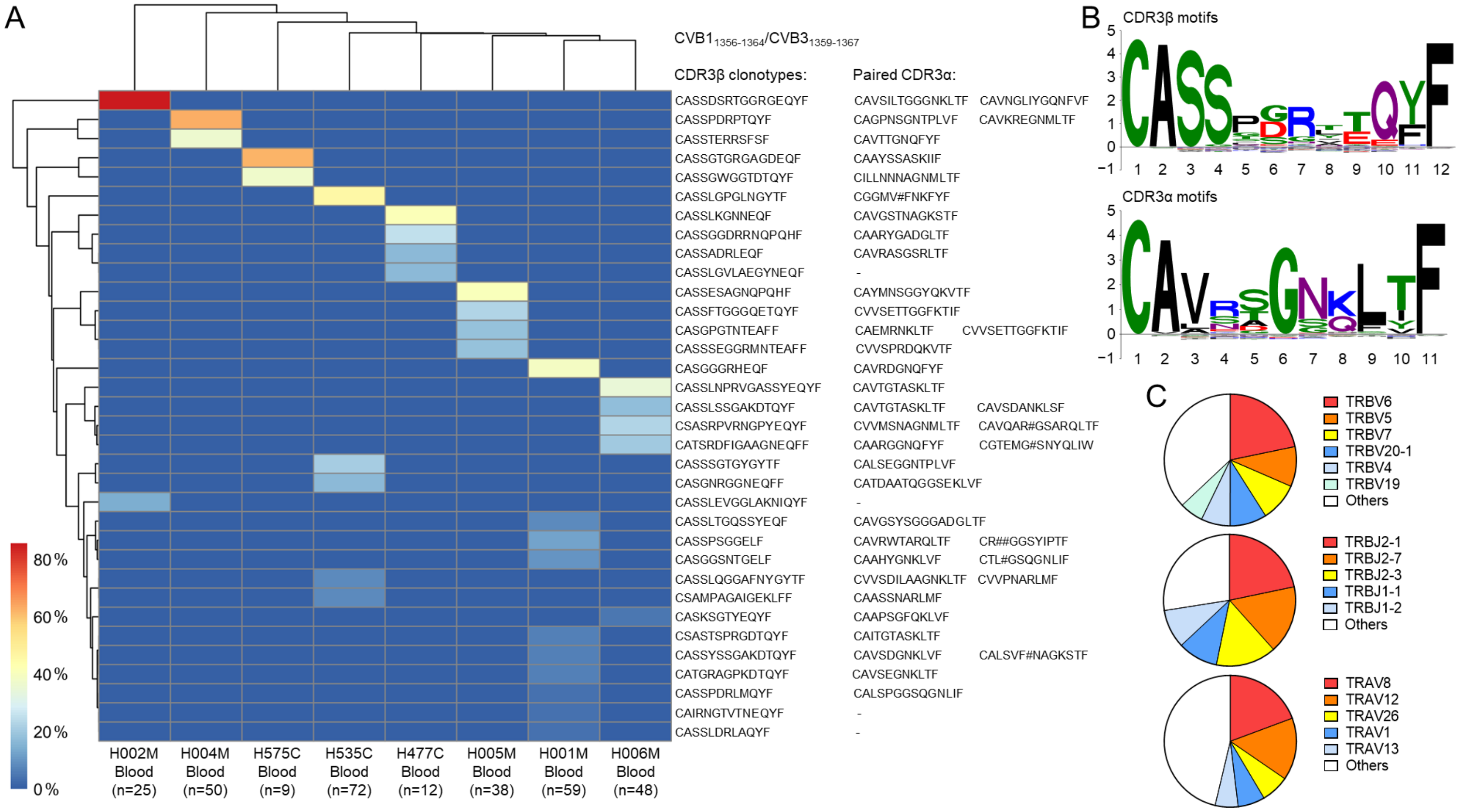

**Figure S6. Expanded TCR CDR3β clonotypes in individual CVB1/CVB3$_{1356\text{-}1364/1359\text{-}1367}$-reactive CD8$^+$ T cells from the blood of living CVB-seropositive healthy donors and common CDR3 motifs. A.** Expanded clonotypes were defined as those found in at least 2 MMr$^+$ cells in the same donor, and are clustered according to frequency (one columns per donor) and sequence similarity (one row per CDR3β clonotype). **B.** Sequence logo plots displaying CDR3β (top) and CDR3α (bottom) motifs among all TCRs sequenced from CVB1/CVB3$_{1356\text{-}1364}$/$_{1359\text{-}1367}$ MMr$^+$CD8$^+$ T cells isolated from tissues and blood. The x-axis shows the CDR3 aa position after MMseqs2 centroid alignment. The y-axis shows the information content, with the size of each aa symbol proportional to its frequency. **C.** Distribution of *TRBV* (top), *TRBJ* (middle) and *TRAV* (bottom) gene usage among 378 TCRs sequenced from CVB1/CVB3$_{1356\text{-}1364}$/$_{1359\text{-}1367}$ MMr$^+$CD8$^+$ T cells isolated from tissues and blood.

A

| | | |
|---|---|---|
| HLA-A2 | CVB1-2-3-4-5-6 1246-1254<br>CVB1-2-4-5-6 271-279 | KLNSSVYSL<br>IVMPYINSV |
| | CMV pp65 495-503<br>Flu MP 58-66 | NLVPMVATV<br>GILGFVFTL |
| | WNV PP 430-438 | SVGGVFTSV |
| HLA-A3 | CVB1 1356-1364 / CVB3 1359-1367<br>CVB1 1132-1140 | KINMPMSVK<br>KILPEVKEK |
| | CMV IE1 99-107<br>Flu NP 265-273 | RIKEHMLKK<br>ILRGSVAHK |
| | WNV PP 3098-3106 | LTYRHKVVK |

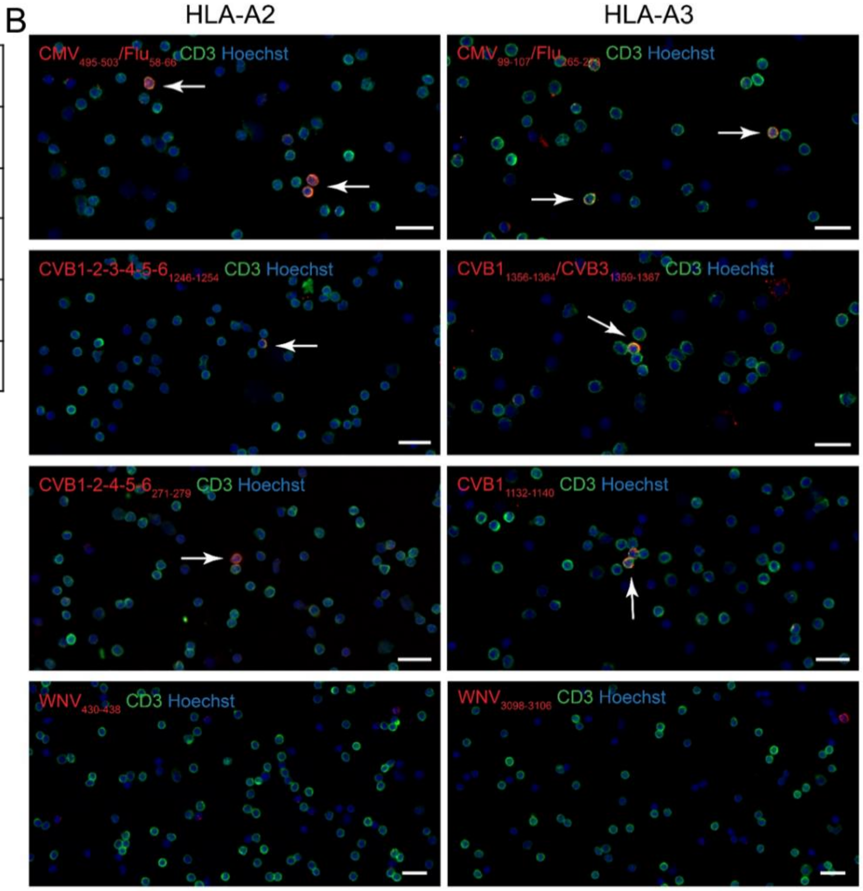

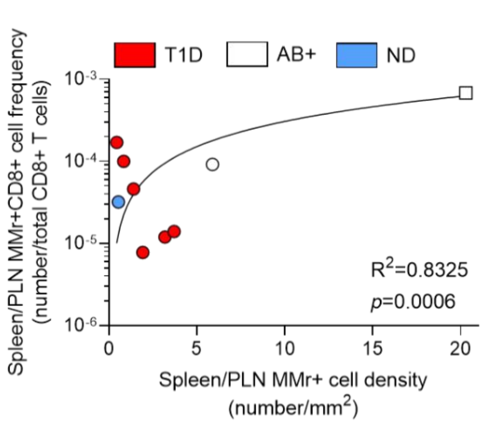

**Figure S7. Validation of *in-situ* MMr staining. A.** MMrs used for *in-situ* staining on human tissues. **B.** Representative immunofluorescence images of isolated $CD8^+$ T cells cytospun on microscope glass slides. Cells were stained with the indicated pooled CMV/Flu, pooled CVB or single WNV peptide-loaded MMrs (red) and CD3 (green). Cell nuclei are stained in blue. White arrows point to $MMr^+CD3^+$ cells. Scale bars: 20 µm. **C.** Correlation between CVB $MMr^+CD8^+$ cell frequencies measured by flow cytometry and CVB $MMr^+CD8^+$ cell densities quantified by tissue immunofluorescence. The samples analyzed by both techniques available for this comparison were all from spleen (round symbols), barring a PLN sample from double-$aAb^+$ case 6197 (square symbol).

A

| | MOI 10 T:β 0.5:1 | MOI 100 T:β 0.5:1 | MOI 300 T:β 0.5:1 | MOI 10 T:β 1:1 | MOI 100 T:β 1:1 | MOI 300 T:β 1:1 | MOI 10 T:β 2:1 | MOI 100 T:β 2:1 | MOI 300 T:β 2:1 |
|---|---|---|---|---|---|---|---|---|---|
| r | +0.93 | +0.70 | +0.97 | +0.16 | +0.92 | +0.97 | −0.75 | −0.96 | −0.85 |
| 95% CI | +0.78<br>+0.98 | +0.26<br>+0.90 | +0.91<br>+0.99 | −0.41<br>+0.65 | +0.76<br>+0.98 | +0.93<br>+0.99 | −0.92<br>−0.37 | −0.99<br>−0.89 | −0.95<br>−0.57 |
| p | <0.0001 | 0.0067 | <0.0001 | 0.5838 | <0.0001 | <0.0001 | 0.0024 | <0.0001 | 0.0002 |

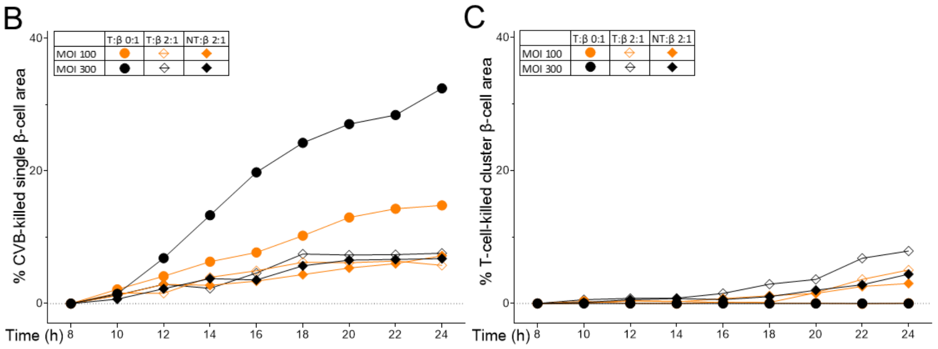

D

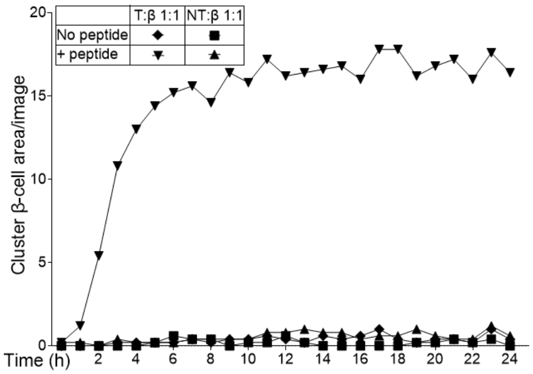

**Figure S8. Correlation between CVB-mediated and T-cell-mediated β-cell killing. A.** r, 95% confidence interval (CI) and *p* values of the correlation between CVB-mediated and T-cell-mediated β-cell killing depicted in Fig. 7D. **B.** Equivalent inhibition of CVB-mediated single β-cell death by both transduced (T) and non-transduced (NT) CD8$^+$ T cells at a 2:1 T:β-cell ratio. **C.** T-cell-mediated cluster β-cell death in the same conditions. **D.** CVB peptide-specific β-cell killing by transduced (T) and non-transduced (NT) CD8$^+$ T cells. No killing is observed with non-transduced T cells.

## A

Structural
HLA-A2-restricted
CVB1 271-279

```
CVB1     - IVMPYINSV
CVB2     - IVMPYINSV
CVB3     - IVMPYTNSV
CVB4     - IVMPYINSV
CVB5     - IVMPYINSV
CVB6     - IVMPYINSV
PV1/2/3  - IVLPYVNSL
```

Non-structural
HLA-A2-restricted
CVB1 1246-1254

```
CVB1     - KLNSSVYSL
CVB2     - KLNSSVYSL
CVB3     - KLNSSVYSL
CVB4     - KLNSSVYSL
CVB5     - KLNSSVYSL
CVB6     - KLNSSVYSL
PV1/2/3  - KENTSTYSL
```

Non-structural
HLA-A3-restricted
CVB1 1132-1140

```
CVB1     - KILPEVKEK
CVB2     - KILPEVKEK
CVB3     - KILPEVREK
CVB4     - KILPEVKEK
CVB5     - KILPEVREK
CVB6     - RILPEVKEK
GAD      - KMFPEVKEK
PV1/2/3  - KIIPQARDK
```

Non-structural
HLA-A3-restricted
CVB1 1356-1364

```
CVB1     - KINMPMSVK
CVB2     - KVNMPMSVK
CVB3     - KINMPMSVK
CVB4     - KINMPMSVK
CVB5     - KINMPMSVK
CVB6     - KINMPMSVK
PV1/2/3  - KLNMAMATE
```

## B

Structural
HLA-A2-restricted
CVB5 169-177

```
CVB1     - YLGRTGYTI
CVB2     - YLGRTGYTI
CVB3     - YLGRTGYTV
CVB4     - YLGRTGYTI
CVB5     - YLGRAGYTV
CVB6     - YLGRTGYTI
PV1/2/3  - YLGRAGYTV
```

Non-structural
HLA-A2-restricted
CVB1 1671-1679

```
CVB1     - RMLMYNFPT
CVB2     - RMLMYNFPT
CVB3     - RMLMYNFPT
CVB4     - RMLMYNFPT
CVB5     - RMLMYNFPT
CVB6     - RMLMYNFPT
PV1/2/3  - RTLMYNFPT
```

Structural
HLA-A3-restricted
CVB5 760-768

```
CVB1     - SCFYDGWTQ
CVB2     - SMFYDGWSE
CVB3     - SNFYDGWSE
CVB4     - TMFYDGWSN
CVB5     - SMFYDGWAK
CVB6     - STFYDGWSD
PV1/2/3  - SHFYDGFAK
```

Non-structural
HLA-A3-restricted
CVB3 1765-1774

```
CVB1     - VLRNGDPRLR
CVB2     - VLRNGDPRLK
CVB3     - VLRSGDPRLK
CVB4     - VLRNGDPRLK
CVB5     - VLRNGDPRLK
CVB6     - VLRNGDPRLK
PV1/2/3  - VLTKSDPRLK
```

**Figure S9. Alignment of the main CD8$^{+}$ T-cell epitopes identified across CVB and Poliovirus serotypes.** Amino-acid differences are shaded in grey. For Poliovirus, the 3 serotypes included in the inactivated poliovirus vaccine were used as reference sequences: serotype 1 (Mahoney; GenBank V01149.1), 2 (MEF1; GenBank AY238473.1) and 3 (Saukett; GenBank KP247597.1). The alignment refers to the serotype with the closest match.

**Movie S1. Kinetics of CVB infection and CVB-eGFP transfer through filopodia in ECN90 β cells.** Real-time imaging of ECN90 β cells infected with CVB-eGFP at MOI 100 and stained with Cytotox Red to visualize dead cells. The movie shows an intact β cell in contact with the filopodia of an infected $eGFP^+$ cell, turning $eGFP^+$ at and subsequently protruding filopodia before dying (Cytotox $Red^+$). Infected cells are labeled in green, dead cells are labeled in red.

**Movie S2. Death kinetics and morphology in CVB-infected ECN90 β cells and definition of a single-cell analysis mask.** β cells (infected at 300 MOI, in the absence of T cells) are labeled in red, dead cells are labeled in green, merged images of dead β cells are labeled in yellow. Contours in magenta red indicate the areas of single-cell death defined by the imaging software based on the preset analysis mask.

**Movie S3. Death kinetics and morphology of ECN90 β cells co-cultured with CVB-reactive $CD8^+$ T-cell transductants and definition of a cell cluster analysis mask.** β cells (pulsed with 1 μM $CVB1_{1356-1364}$ peptide, without infection) are labeled in red, dead cells are labeled in green, merged images of dead β cells are labeled in yellow. Contours in magenta red indicate the areas of clustered cell death defined by the imaging software based on the preset analysis mask.

**Movie S4A-B. Mask analyses for counting of single-cell and cluster cell death areas of CVB-infected ECN90 β cells co-cultured with CVB-reactive $CD8^+$ T-cell transductants.** β cells (infected at 300 MOI and co-cultured at a 1:1 T:β-cell ratio) are labeled in red, dead cells are labeled in green, merged images of dead β cells are labeled in yellow. The analysis of the same field with the defined single-cell (A) and cluster cell analysis mask (B) is shown, with contours in magenta red indicating the areas counted with each mask.
